# ECD co-operates with ERBB2 to promote tumorigenesis through upregulation of unfolded protein response and glycolysis

**DOI:** 10.1101/2025.01.28.635284

**Authors:** Benjamin B. Kennedy, Mohsin Raza, Sameer Mirza, Asher Rajkumar Rajan, Farshid Oruji, Matthew M. Storck, Subodh M. Lele, Timothy E. Reznicek, Lusheng Li, M. Jordan Rowley, Shibiao Wan, Bhopal C. Mohapatra, Hamid Band, Vimla Band

**Affiliations:** Department of Genetics, Cell Biology, and Anatomy, College of Medicine, University of Nebraska Medical Center, Omaha, Nebraska; Department of Pathology, Microbiology, and Immunology, University of Nebraska Medical Center, Omaha, Nebraska; Fred and Pamela Buffett Cancer Center, University of Nebraska Medical Center, Omaha, Nebraska; Eppley Institute for Research in Cancer and Allied Diseases, University of Nebraska Medical Center, Omaha, Nebraska

**Keywords:** ECD, ERBB2, transgenic mice, mammary tumors, hMECs, UPR, glycolysis and mRNA stability

## Abstract

The ecdysoneless (ECD) mRNA and protein are overexpressed in breast cancer (BC), and its overexpression correlates with poor prognosis and short patient survival, particularly in ERBB2/HER2-positive BC. This study investigates the co-operative oncogenic mechanism of ECD and ERBB2 by deriving transgenic mice overexpressing ECD and/or ERBB2 (huHER2) in mammary epithelium under MMTV promoter, as well as human mammary immortal epithelial cell lines (hMECs) overexpressing ECD and/or ERBB2. While *huHER2*Tg mice developed more homogenous solid nodular carcinomas, double transgenic mice (*ECD;huHER2*Tg) developed heterogenous and histologically aggressive mammary tumors with basal-like phenotype and epithelial mesenchymal transition (EMT) features, like *ECD*Tg tumors, resembling more to patient tumors. Importantly, transcriptomic profile of *ECD;huHER2*Tg tumors revealed upregulation of two major oncogenic pathways, unfolded protein response (UPR) and glycolysis. Similarly, hMECs expressing both ECD and ERBB2 as compared to single gene expressing cells showed increase in oncogenic traits, and RNA-seq analysis showed a significant upregulation of glycolysis and UPR pathways. ECD is an RNA binding protein, and directly associates with three key glycolytic enzymes (*LDHA*, *PKM2* and *HK2*) and mRNA of a major UPR regulated gene GRP78, that results in increased mRNA stability. Lastly, we show an increase in glucose uptake and enhanced glycolytic rate in ECD+ERBB2-overexpressing cells as compared to ECD− or ERBB2-overexpressing hMECs. Taken together, our findings support a co-operative role of ECD and ERBB2 in oncogenesis by enhancing two major oncogenic pathways, UPR and glycolysis.

**Significance:** This study provides mechanistic insights that overexpression of ECD in ERBB2+ breast cancer patients correlates with shorter patient survival, by identifying direct ECD binding to mRNAs for UPR and glycolysis pathways.

## INTRODUCTION

Among the subtypes of breast cancer (BC) that with ERBB2/HER2 overexpression, representing about 15-20% of all cases, is associated with an inherent aggressive clinical phenotype and poor prognosis (1, 2). While the outcomes of ERBB2+ BC patients have dramatically improved with the introduction of targeted therapy with humanized anti-ErbB2 antibodies, Trastuzumab (Herceptin) or Trastuzumab plus Pertuzumab + (PERJETA), in combination with chemotherapy, these treatments are non-curative in recurrent and metastatic disease (3). Recurrence and metastasis are attributed to alternate signaling pathways, such as those impinging on the activation of the PI3K/AKT/mTOR axis, and metabolic reprogramming (4). A major adaptive metabolic change in tumor cells is the preferential utilization of aerobic glycolysis, also known as the Warburg effect, to reallocate glycolytic intermediates for biosynthetic needs and to reduce the generation of reactive oxygen species from oxidative phosphorylation (5). Pyruvate kinase muscle isozyme M2 (PKM2, catalyzes the conversion of phosphoenolpyruvate and ADP to pyruvate and ATP), Lactate dehydrogenase A (LDHA, is required to maintain glycolysis and ATP production in the absence of sufficient oxygen by regenerating NAD ^+^ form NADH) and Hexokinase 2 (HK2, catalyzes the phosphorylation of glucose, the rate-limiting first step of glycolysis) are key among the enzymes involved in redirecting glycolytic intermediates into the anabolic pathway (6). Consistent with their key roles, inhibition of HK2, LDHA or PKM2 has been shown to sensitize tumor cells to antitumor effects of therapeutic agents (7). 2-Deoxy glucose (2-DG), an HK2 inhibitor, has been shown to sensitize cancer cells to chemotherapy or radiotherapy and is being investigated in pre-clinical trials (8). Thus, novel adaptive mechanisms that promote the glycolytic switch are likely to play key roles in promoting ERBB2+ BC oncogenesis.

In addition, increased biosynthetic and metabolic demands of tumor cells necessitate increased synthesis of a broad array of cellular proteins, including secretory and transmembrane proteins synthesized in the endoplasmic reticulum (ER) (9, 10). For ERBB2+ BC, the driver oncogene itself is an ER-synthesized transmembrane protein, and its overexpression further increases the load of unfolded proteins in the endoplasmic reticulum (ER stress) (11). The ensuing unfolded protein response (UPR) mediated by ER-resident sensors of misfolded protein load triggers homeostatic pathways that shut down new mRNA translation and promote mRNA degradation to reduce the load of unfolded proteins while concurrently, activating transcription of genes whose products promote ER protein folding and enhance ER-associated degradation of misfolded proteins. Activation of UPR is associated with a block of cell cycle progression (12); the UPR activation also commits cells with unresolved ER-stress to apoptotic death (13). Indeed, ERBB2+ BC show increased expression of genes encoding proteins involved in the UPR and ERAD pathways (11, 14). However, to counteract the negative impact of the overactive UPR, tumor cells must adapt to promote its beneficial aspects while counteracting its anti-proliferative and pro-apoptotic impact. Thus, defining such biological adaptations will help elucidate the pathways that promote oncogenesis in ERBB2-positive breast cancers.

Work from our laboratory and others has helped establish that human ECD (ecdysoneless, a Drosophila homolog), an evolutionarily conserved protein is a positive regulator of cell cycle progression (15) and cell survival (16). *ECD* mRNA is overexpressed in more than 25% of BC patients in TCGA cohort and more than 10% in METABRIC cohort (17). However, ECD protein is overexpressed in more than 85% of ERBB2+ BC patient samples (18) and such overexpression correlates with poor prognostic markers and shorter patient survival in ERBB2+ BC (17, 18). ECD knockdown in ERBB2-positive BC cell lines resulted in decreased oncogenic traits (17), supporting a biological role of ECD overexpression in sustaining ERBB2-driven oncogenesis. However, the experimental proof of ECD promoting ERBB2-driven oncogenesis and the mechanistic insights into how ECD co-operates with ERBB2 to drive oncogenesis, are lacking.

To address these questions, we generated mammary epithelium-targeted human ECD and ERBB2 double transgenic mice (*ECD;huHER2*Tg), as well as ECD and/or ERBB2 overexpressing immortalized human mammary epithelial cells (hMECs) as human cell models. To generate double transgenic mice, we crossed the MMTV-driven *ECD*Tg mice, a majority (about 70%) of which develop mammary hyperplasia with 33% exhibiting heterogeneous mammary tumors (19), with MMTV-driven human ERBB2 transgenic (huHER2) mice (20). We show that the *ECD; huHER2*Tg mice exhibit heterogenous, histologically aggressive and highly proliferative tumors like *ECD*Tg mice. Similarly, ECD+ERBB2-overexpressing hMECs exhibited significantly more oncogenic traits as compared to hMECs expressing single gene. Significantly, *ECD;huHER2*Tg mice tumors and ECD+ERBB2-overexpressing hMECs exhibited increased mRNA levels of glycolytic enzymes, and glucose-regulated protein 78 (GRP78), a key player in UPR. Given our recent findings that ECD directly binds to cellular mRNAs (21), we analyzed and demonstrate here that ECD binds to mRNAs of key glycolytic enzymes, LDHA, PKM2 and HK2 as well as GRP78, and this interaction results in enhanced mRNA stability. Cell based analyses demonstrated ECD overexpression-dependent enhancement of glucose uptake and glycolytic rates in hMECs expressing both ECD and ERBB2. Taken together, our results establish that ECD overexpression co-operates with ERBB2 to drive more aggressive oncogenesis through regulation of glycolysis and UPR.

## MATERIALS AND METHODS

### Generation of mammary-epithelium targeted ECD/ERBB2 (HER2) double-transgenic mice

The *MMTV.huHER2* mice with a human ERBB2 (HER2) transgene under an MMTV promoter (20) were obtained from Genentech Inc., and the Tet-off human ECD transgenic mice, *Tet(O)-ECD; MMTV-tTA* (referred to as *ECDTg*), generated previously (19) were bred to generate the *Tet(O)-ECD; MMTV-tTA; MMTV.huHER2* double transgenic (DTg) mice. The presence of all alleles was verified by tail clip DNA PCR genotyping using the primer sequences described previously for *Tet(O)-ECD* and *MMTV-tTA* (19) and *MMTV-huHER2* (20). Transgenic mice were maintained under specific pathogen-free conditions at the Center for Comparative Medicine of the University of Nebraska Medical Center. All animal studies were pre-approved by the Institutional Animal Care and Use Committee (IACUC) under protocol number 20-008-04-FC. All experimental analyses were carried out using female mice as our focus is on breast cancer, which predominantly affects women.

### Immunohistochemistry

Immunohistochemistry (IHC) of tumor sections was carried out as described previously (19). Briefly, a portion of the excised mammary tumor was fixed in 10% neutral buffered formalin (cat#316-155; PROTOCOL), embedded in paraffin and 5-mm thick sections prepared. The sections were deparaffinized in xylene, rehydrated in descending alcohols, and treated in a digital pressure cooker containing citrate buffer (pH 6.0; DakoCytomation, S1699) for antigen retrieval. Endogenous peroxidase activity was blocked by incubation in 3% hydrogen peroxide for 20 min. The sections were rinsed in phosphate buffered saline (PBS) and incubated for 15 min in protein block buffer (DakoCytomation #0909). The sections were then stained with the respective primary antibodies overnight at 4^0^C. After rinsing in Tris-buffered saline with 0.1 % Tween-20 (cat#1610781, BIO-RAD) (TBST), the sections were incubated for 15 min with an anti-mouse or anti-rabbit secondary antibody conjugated to a dextran-labeled polymer with horseradish peroxidase (HRP) (DakoCytomation K4007), followed by incubation in DAB solution (DakoCytomation DAB # K4007, Solution 3a, b) for 3 to 7 min. The sections were counter stained in hematoxylin and mounted under cover glasses using a xylene-based mounting medium using standard protocols.

### TERT-immortalized human mammary epithelial cell (hMEC) lines 76NTERT and 70NTERT and generation of ECD, ERBB2 or ECD+ERBB2-overexpressing hMEC cell lines

The hTERT immortalized hMECs, 76NTERT and 70NTERT, were cultured at 37°C under 5% CO_2_ in growth factor and serum containing DFCI-1 medium (22, 23). Growth factor and serum-depleted DFCI-3 medium (22, 23) was used for specific experiments as indicated. Full-length human ECD and ERBB2 were cloned into pMSCV-hygromycin and pMSCV-puromycin retroviral vectors (17, 24) respectively. The empty vectors or ECD/ERBB2 constructs were transiently transfected into Plat-GP packaging cell line (cat# RV-103, Cell BIOLABS) grown in DMEM with 10% FBS (fetal bovine serum) and retroviral supernatants were used for transduction of cells in the presence of 10 μg/ml Polybrene (cat# TR-1003-G, Sigma-Aldrich). Transduced cells were cultured in media with 5 μg/ml hygromycin B (cat# 10687010, Thermo Fisher Scientific) and/or 1 μg/mL puromycin (cat# P8833, Sigma-Aldrich). Cell lines were regularly tested for mycoplasma and continuously cultured for no more than 3 months.

### Antibodies, cell lysate preparation and western blotting

The antibodies used were: Anti-ECD mouse monoclonal antibody (18); anti-ECD (cat# 10192-1-AP, Proteintech); Anti-ERBB2 (cat# 2242, Cell Signaling Technology); Anti-b-actin (cat# AC-15, Thermo Fisher Scientific); and anti-Rabbit IgG antibody (cat# 7074P2, Cell Signaling Technology). Cell lysates were prepared in RIPA lysis buffer (cat# 89901, Thermo Fisher Scientific) supplemented with phosphatase and protease inhibitor cocktails (cat# P8340 & P0001, Sigma-Aldrich) and quantified using the BCA method (cat#23226, Thermo Fisher Scientific). Acrylamide gel-resolved proteins were transferred to PVDF membrane (cat# IPVH00010, Millipore). The membranes were blocked in 5% bovine serum albumin for one hour at room temp, incubated with the primary antibody diluted in TBST. Following three times in TBST (cat#J77600.K8, Thermo Fisher Scientific), membranes were incubated in HRP conjugated secondary antibodies (Zymed Laboratories), washed in TBST three times and signals were detected using PerkinElmer Western Lighting Plus-ECL solutions (cat# 50-904-9326, Thermo Fisher Scientific).

### CellTiter-Glo proliferation assay

96-well Optical plates (Thermo Scientific 165305) were seeded with 1,000 cells per well in replicates of 6 in either complete DFCI or DFCI-3 starvation (23) medium and cultured at 37°C with 5% CO_2_. Viable cells at the indicated time points were quantified using the Promega CellTiter-Glo kit (G7571) according to the manufacturer’s protocol (25).

### Glucose Uptake-Glo assay

Cells were seeded as for the proliferation assay except for 25,000 cells per well. 24 hours after plating, the cells were washed twice with PBS and cultured in glucose-free DMEM media (cat#11966025, Thermo Fisher Scientific) for one hour cells glucose starvation. The Promega Glucose Uptake-Glo (J1341) assay was then performed according to the manufacturer’s protocol (26). Parallel CellTiter-Glo assays were used for normalization of cell numbers.

### Seahorse XF Glycolytic Rate Assay

The Oxygen Consumption Rate (OCR), Extracellular Acidification Rate (ECAR) and Photon Efflux rate (PER) were measured using a Seahorse XFe Glycolytic rate assay Kit (cat #103344-100, Agilent Technologies) according to the manufacturer’s protocol (27). Briefly, 3.5 × 10^4^ 76NTERT or 70NTERT gene transduced cells were seeded per well in 96-well seahorse XFe cell culture plates and incubated overnight. Cells were washed and incubated with XF assay medium supplemented with 1 mM pyruvate (cat# S8636, Sigma-Aldrich), 2 mM L-glutamine (cat#25030081, Invitrogen), 10 mM glucose (cat#A24940-01, Gibco) for 1 h at 37°C in a CO_2_ free incubator. The OCR, ECAR and PER estimations were performed as per the manufacturer’s instructions. Basal OCR and ECAR was assessed in response to rotenone/antimycin A (0.5 μM), and 2-deoxy-D-glucose (2-DG; 50 mM). ATP rate was estimated in response to oligomycin (1.5 μM) and rotenone/antimycin A (0.5 μM). The readings were normalized to the respective protein concentrations. Basal glycolysis was calculated from non-glycolytic acidification (background) ECAR/PER. Glycolytic reserve (Compensatory) was calculated as the difference between glycolytic capacity and basal glycolysis.

### Anchorage independent growth assays

20,000 cells suspended in 2 mL 0.3% agarose in DFCI-1 medium were plated per well in 6-well plates on top of pre-solidified 2 mL layer of 0.6% agarose in DFCI-1. 2 mL of fresh DFCI-1 medium was changed every 72 hours for 3 weeks, and colonies stained with 0.05% crystal violet were imaged in six random fields per well at 4x and 20x magnifications. The average number of colonies and colony size were quantified using the ImageJ software.

### Transwell migration and invasion assays

Regularly fed cell cultures subjected to serum and growth factor starvation by culturing in DFCI-3 medium for 24 hours were trypsinized and plated at 20,000 cells per well in DFCI-3 medium in upper chambers of transwell migration (Corning BioCoat 8.0 μm #354578) or invasion (BioCoat Matrigel 8.0 μm #354480) transwell plates. After two hours of incubation at 37^0^C, DFCI-1 medium was added as a chemoattractant in the bottom chambers. After 24 hours of incubation, the cells in upper chambers were removed with cotton swabs and the migrated cells were rinsed in PBS, fixed in ice-cold methanol, stained with crystal violet, and rinsed again in PBS. Cells in three random fields per well were imaged at 10x magnification and quantified using the ImageJ software.

### Mammosphere Assay

Mammosphere assay was carried out as described (28). Briefly, 3,000 cells were plated in 96-well ultra-low attachment plates (Corning No. 3474) in mammosphere medium. The mammospheres were imaged five days after plating at 40x magnification and quantified using the ImageJ software (NIH), using the 50 μm diameter as the size cut-off. All experiments were performed in 6 replicates and repeated three times.

### Matrigel Organoid Assay

Single cell suspensions of cells were prepared in the DFCI-1 complete medium containing 4% Matrigel (cat#356231, Corning) and seeded at 5,000 cells/well on the top of a base layer of 100% Matrigel in eight-well Nunc chamber slides (Nunc Lab-Tek Chamber slide, cat#C7182, MilliporeSigma). The cells were allowed to form organoids for 5 days and then imaged, with quantification using the ImageJ software with a size cut-off of 50 μm in diameter.

### RNA isolation and real time quantitative PCR

Total RNA was isolated from mammary tumor samples or cultured cells using the Trizol reagent (cat#15596018, Ambion), according to the manufacturer’s protocol. The RNA was reverse transcribed into cDNA using the BioRad iScript gDNA Clear cDNA Synthesis Kit (cat#1725035, Bio-Rad). For real-time (RT)-qPCR, triplicate samples were analyzed using Applied Biosystems SYBR Green master mix (cat# 4309155) and 1 μg cDNA diluted in nuclease-free water. Relative fold change in RT-qPCR products was calculated using the ΔΔCt method and expressed relative to β-actin as a housekeeping gene. Primers were purchased from Sigma-Aldrich, Eurofins, or Invitrogen. All qPCR reactions were performed using an Applied Biosystems QuantStudio3 system according to SYBR Green reaction parameters as defined by the manufacturer. The RT-qPCR primers for human and mouse mRNAs are listed in **supplementary Tables S1 and S2**.

### RNA Immunoprecipitation (RIP) Assay

Total RNA was isolated from 76NTERT cells using the standard RIP protocol without crosslinking (29). Briefly, 76NTERT cells were cultured in complete DFCI medium, and lysates prepared in cold lysis buffer (50 mM HEPES, Triton-X 10%, NP40 0.5%) were frozen at −80°C overnight before thawing on ice. For the RIP, 2.5 ug/mL of anti-ECD (cat # 10192-1-AP, Proteintech) or anti-Rabbit IgG antibody (cat # 7074P2, Cell Signaling Technology) bound on protein A/G immuno-magnetic beads were mixed with cell lysates (1 mg/mL) to immunoprecipitate (IP) the ECD-mRNA complexes. The RNA in the precipitated protein-RNA complexes was purified using the standard TRIzol reagent per the manufacturer’s instructions. DNA free cDNA was prepared using the Bio-Rad iscript gDNA clear cDNA synthesis Kit (cat #1725035). and subjected to qRT-PCR to measure the levels of mRNAs (*HK2, LDHA* and *PKM2* and *HSPA5*).

### Actinomycin D RNA Stability Assay

RNA stability experiments were carried out as described previously (17, 19). 76NTERT transductants cultured in serum-free DFCI-3 medium for 72 h were treated with 5 μg/mL actinomycin D (Sigma Aldrich A1410) to inhibit the global mRNA transcription. Cells were harvested at the indicated time points to isolate RNA using the Trizol reagent (cat#15596018, Ambion). Subsequent cDNA synthesis and qRT-PCR analyses were as described above. The half-life for mRNA transcript decay was calculated using GraphPad Prism software.

### Transcriptome profiling of tumors by RNA-seq analysis

Total RNA was isolated using the Trizol reagents from a central piece of excised mammary tumors, as described (19). The RNA preparations were treated with DNase I and further cleaned using the Qiagen RNeasy clean up and micro kit, as per the manufacturer’s protocols. The purity of RNA was assessed on a Bioanalyzer at the UNMC Next Generation Sequencing Facility. 1-2 mg of cleaned RNA samples was used to generate RNA-seq libraries using the TruSeq RNA Library Prep Kit v2 (Illumina) following the manufacturer’s protocols and sequenced using the 2 x 100 bases paired-end protocol on a NovaSeq 6000 instrument (Illumina). The NGS short reads from the RNA-seq experiments were downloaded from the NovaSeq 6000 server in FASTQ format. Raw FASTQ reads were trimmed using the *Trim Galore* v0.6.5 (30) to remove the adapters, terminal unknown bases (Ns), and low-quality regions (Phred score < 33). The reads that passed the quality control were mapped to the mouse genome assembly (mm10) using STAR v2.7 (31) and the gene level values were quantified using RSEM v1.3 (32) based on GENCODE annotation (v35). For differentially expressed gene (DEG) analysis between the experimental conditions, the expected read counts from tumor samples were used. Genes with low read counts were filtered out using a CPM cutoff threshold equivalent to a count of 10 reads. Normalization factors were then calculated using the TMM method. The counts were further normalized using the voom and normalized counts were analyzed using the lmFit and eBayes functions from the limma R package v3.54.2 (33). Genes with adjusted p-value < 0.05, and absolute fold change > 2 were considered as significantly up- and down-regulated genes. Heatmaps were generated using R Bioconductor package heatmap. Gene Set Enrichment Analysis (GSEA) v4.3.2 (34) was performed between the experimental conditions to analyze signaling pathways using MSigDB gene sets.

### RNA Sequencing and Analysis of 76NERT cell line transfectants

76NTERT cells expressing vector, ECD, ERBB2, and ECD+ERBB2 were cultured in triplicates in DFCI-3 starvation medium for 72 h. Following total RNA isolation and Bioanalyzer analysis for purity as above, the RNA-seq libraries were prepared with the Illumina TruSeq Stranded RNA Library Preparation Kit. The RNA-seq reads underwent processing via the nf-core RNA-seq pipeline (version 3.9) (35), accessed through Nextflow (version 22.10.3 (36), and executed with Singularity (37) to ensure reproducibility and portability. Utilizing paired-end sequencing reads, alignment to the human reference genome (GRCh38) was achieved using the Spliced Transcripts Alignment to a Reference (STAR) software (version 2.7.11a), integrating Ensembl gene annotations for precise mapping. Transcript abundance quantification was conducted using StringTie2 (version 1.3.6), with transcripts per million (TPM) normalization for inter-sample comparisons as described previously (25). Differential expression analysis was performed with DESeq2, identifying significantly differentially expressed genes based on adjusted p-value < 0.05 and a minimum two-fold change in expression (25). Functional interpretation of the differential expression results utilized the Gene Set Enrichment Analysis (GSEA) software from the Broad Institute (34), generating enrichment plots to visualize gene set distribution to identify pathways significantly associated with observed transcriptional changes.

### Statistical analyses

Each assay was carried out three times, with at a minimum three technical replicates within each experiment. Means and standard error of the mean (SEM) are indicated in figure legends. Statistical significance of differences in proliferation were determined using two-way ANOVA with a significant p-value cutoff of 0.05. All other statistical significances were calculated using unpaired two-tailed student’s t-tests with Welch’s correction with a significant p-value cut off of 0.05. Statistically significant differences between experimental groups are indicated in figures using stars (*p<0.05, **p<0.01, ***p<0.001).

### Data Availability

The RNA-seq files are available from the NCBI Gene Expression Omnibus (GEO) under the accession number GSE287075.

### Human and animal subjects

No human subjects were used in this study. Use of mice in this study was pre-approved by the Institutional Animal Care and Use Committee (IACUC) under protocol number 20-008-04-FC that follows Federal and State guidelines.

## RESULTS

### Crossing of *ECD*Tg mice with *huHER2*Tg mice resulted in aggressive and heterogenous mammary tumors

Our previous studies that ECD and ERBB2 co-overexpression is a poor prognostic marker, and it correlates with shorter survival in ERBB2-positive BC patients as compared to patients overexpressing ECD or ERBB2 alone (17, 18) suggesting oncogenic cooperation between ECD and ERBB2. To experimentally address such cooperativity in an *in vivo* setting, we generated mammary epithelium-targeted human *ECD;huHER2* double Tg mice (see method) and compared them with single *ECD*Tg and *huHER2*Tg mice for mammary tumor development over time. The latency of tumor development in double-Tg mice was comparable to that in *huHER2*Tg mice and the percentage of mice that developed tumors were comparable to that of *huHER2*Tg and *ECD*Tg mice **(Supplementary Table S3)**.

IHC analyses using anti-ECD and anti-ERBB2 antibodies confirmed the expected expression of ECD and ERBB2 protein according to the genotype **(Fig. 1A &B)**. Histologically, the *ECD*;*huHER2*Tg (refereed as double Tg) tumors exhibited a heterogeneous morphology with distinct histologic subtypes (**Fig. 1C**), which was like *ECD*Tg alone. Of the 8 tumors observed in double Tg (DTg) mice, three adenosquamous carcinoma type, 3 showed solid nodular carcinoma morphology with epithelial cells arranged in sheet-like structures, and 2 were spindle cell carcinoma with EMT features (**Fig. 1C**). In contrast, all the tumors observed in *huHER2*Tg mice displayed a solid nodular morphology, like previous reports (38) (Supplementary **Table-S3; Fig. 1C**). Notably, three independent *ECD*Tg tumors of adenosquamous carcinoma, spindle cell carcinoma and solid nodular carcinomas were comparable with tumors seen in DTg. Notably, the *ECD*;*huHER2*Tg tumors histologically resemble the tumors that develop in *ECD*Tg mice, which exhibit heterogeneous morphologies (19) similar to other genetically engineered mouse models of BC, such as *C-MYC* and *WNT* (39). Given that HER2-positive patients exhibit tumor heterogeneity, our DTg mice derived tumors resemble phenotype seen in patients (40) rather than *huHER2*Tg mice.

**Fig. 1:**
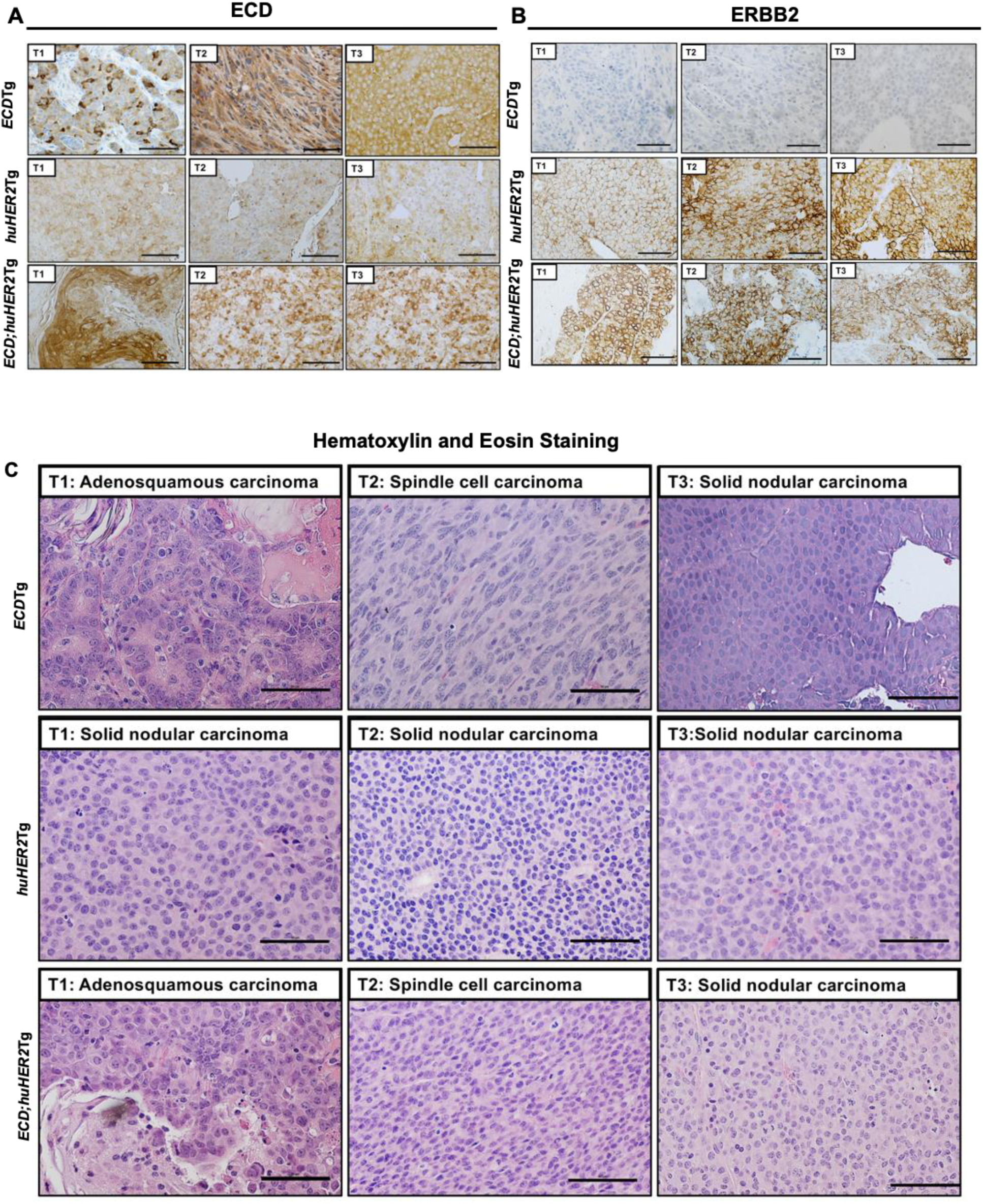
Histopathology of tumors from *ECD*Tg*, huHER2*Tg*, and ECD;huHER2*Tg show tumors from double transgenic mice are heterogenous, like *ECD*Tg tumors. **(A & B)** Deparaffinized tumor sections were stained with indicated antibodies. Representative immunohistochemistry staining images of ECD and ERBB2 are shown. **(C)** Representative images of hematoxylin and eosin-stained sections. Three independent tumors from each Tg mice are shown. Magnification of images are 400x and Scale bar, 50 µm.

Given the substantial increase in the histopathological heterogeneity seen in H&E sections of the DTg tumors, we carried out IHC analyses of a few tumors associated markers. Notably, *ECD;huHER2*Tg tumors exhibited higher Ki67 marker staining as compared to *huHER2*Tg tumors indicating a higher degree of proliferation in DTg tumors (**Fig. 2A & B**). *ECD;huHER2*Tg tumors also exhibited strong cytokeratin (CK)14 positivity, akin to that with *ECDTg* tumors (**Fig. 2C**) (19). Furthermore, the DTg mice tumors showed increased nuclear p63, a known basal (squamous epithelial) cell marker (39) (**Fig. 2D**). Importantly, both *huHER2*Tg and DTg tumors express luminal marker, cytokeratin (CK)18 as expected, like *huHER2*Tg tumors that are known to express uniform luminal cell type (38) (**Fig. 2E**). However, DTg tumors express both basal and luminal markers like adenosquamous type of *ECD*Tg tumors indicating high degree of inter-tumoral heterogeneity with mixed cell lineages (**Fig. 2E**). Given the increased basal expression in *ECD*Tg tumors (19), we also examined the expression of EMT (epithelial to mesenchymal transition) markers. The DTg tumors showed higher expression of vimentin and TWIST and weak E-cadherin staining; in contrast, *huHER2*Tg tumors showed strong E-cadherin staining, moderate vimentin staining and negative staining of TWIST. *ECD*Tg tumors display strong E-cadherin staining, strong vimentin staining and moderate TWIST staining (**Fig. 2F-H**), as reported previously for the *huHER2*Tg tumors (20, 38, 41) and for the *ECD*Tg tumors (19). Overall, our analyses of double vs. single Tg mouse models indicate that crossing of *ECD*Tg with huHER2Tg resembles latency and proportion of mice with *huHER2*Tg mice, the tumor heterogeneity was like *ECD*Tg, suggesting *in vivo* ECD may contribute to heterogenous aggressive tumor phenotypes in ERBB2/HER2-positive tumors in patients.

**Fig. 2:**
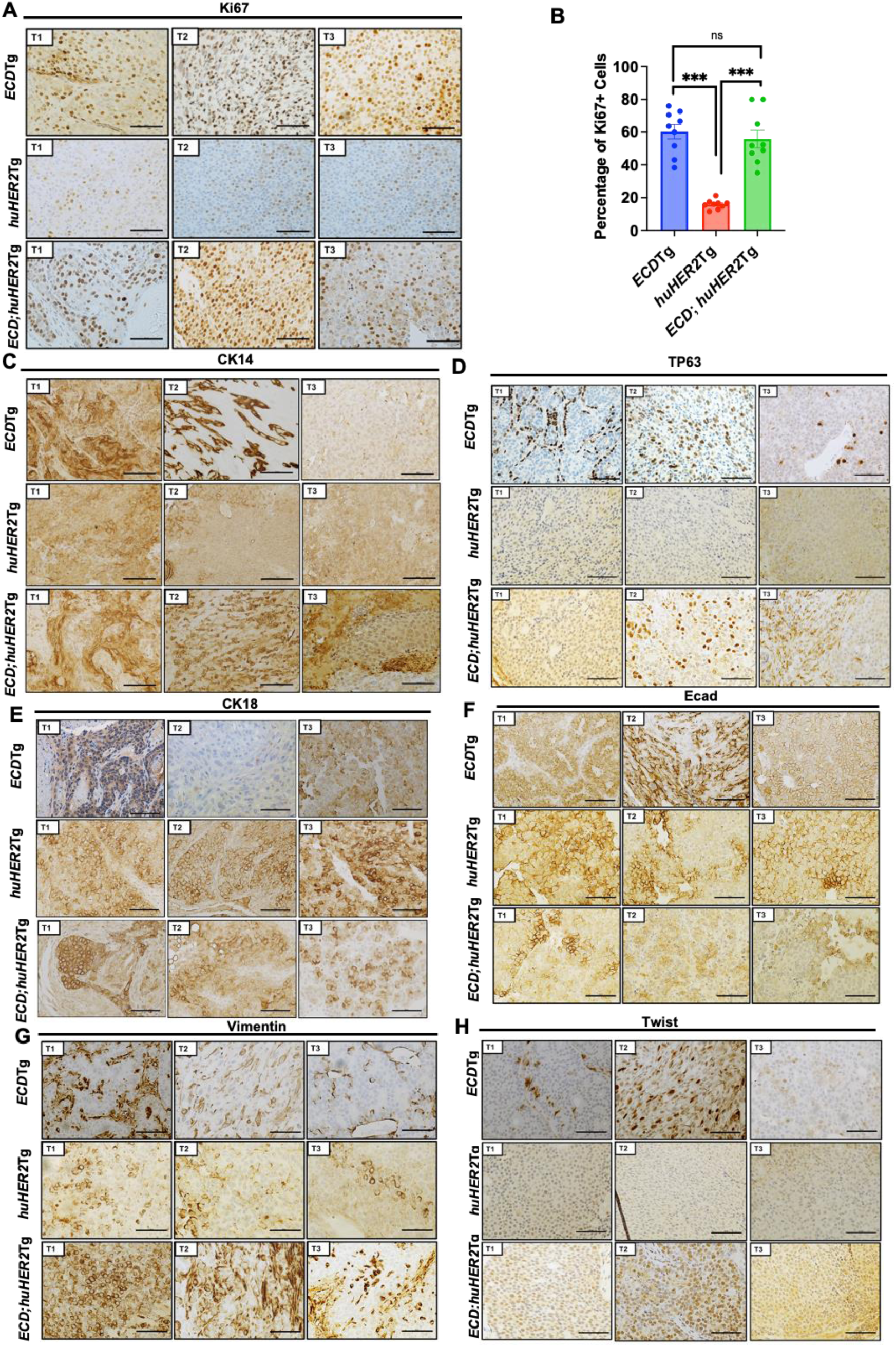
Immunohistochemistry of *ECD*Tg, *huHER2Tg* and *ECD;huHER2*Tg tumors using indicated markers confirms histological heterogeneity. **(A, C-H)** Deparaffinized tumor sections from three Tg tumors were stained with indicated antibodies, proliferative marker Ki-67 **(A).** Quantitation of proliferative marker Ki-67 in independent tumors from each group, represented as percentage of Ki-67+ cells. Data represents mean ± SEM with two-tailed unpaired t test. n = 3; ns, P > 0.05; ***, P < 0.001 **(B)**. Squamous epithelial marker, cytokeratin 14 (CK14) **(C),** basal marker: p63 **(D)**, luminal marker cytokeratin 18 (CK18) **(E)**, epithelial marker: E-cadherin **(F)**, and epithelial to mesenchymal transition markers: vimentin (**G**) and twist **(H)**. Images are taken in 400x magnification, Scale bar represents 50 µm.

### Transcriptome profiling identifies upregulation of unfolded protein response (UPR) and glycolysis pathways in *ECD;huHER2*Tg tumors

To identify the molecular pathways altered upon ECD and ERBB2 co-overexpression, we carried out RNA sequencing analysis of three tumors from each of *ECDTg, huHER2*Tg, and *ECD;huHER2*Tg DTg mice. Principal component analysis (PCA) showed distinct clustering of the tumor groups (**Fig. 3A**). Differential gene expression (DEG) analysis showed an upregulation of 1,126 genes and downregulation of 3,720 genes in *ECD;huHER2*Tg DTg tumors compared to *huHER2* tumors based on a fold change >2 and false discovery rate (FDR) of <0.05. Compared to *ECD*Tg tumors, the *ECD;huHER2*Tg DTg tumors exhibited 5,037 genes upregulation ad 7,430 genes downregulation. Gene set enrichment analysis (GSEA) using the hallmark gene sets revealed a significant upregulation of unfolded protein response (UPR), protein secretion, glycolysis, oxidative phosphorylation, mTORC1 signaling, and PI3K-AKT-mTOR signaling in *ECD*; *huHER2*Tg DTg tumors as compared to *huHER2*Tg (**Fig. 3B**) or *ECD*Tg (**Supplementary Fig. S1A**) tumors. These are well known upregulated pathways in aggressive breast tumors (42). In addition, E2F targets, G2/M check point, MYC targets V1 and V2, and hallmark signaling pathways of proliferating and metastatic tumors were upregulated in ECD;*huHER2*Tg DTg tumors (**Fig. 3B-F**). Importantly, key genes in the UPR (*HSPA5*, *HSP90B1*, *XBP1*, *ATF4*) and glycolysis (*ALDOA, LDHA, PKM, PGK1*) pathways were upregulated in double Tg compared to single Tg tumors (**Fig. 3D, 3F and Supplementary Fig. S1B-I**).

**Fig. 3:**
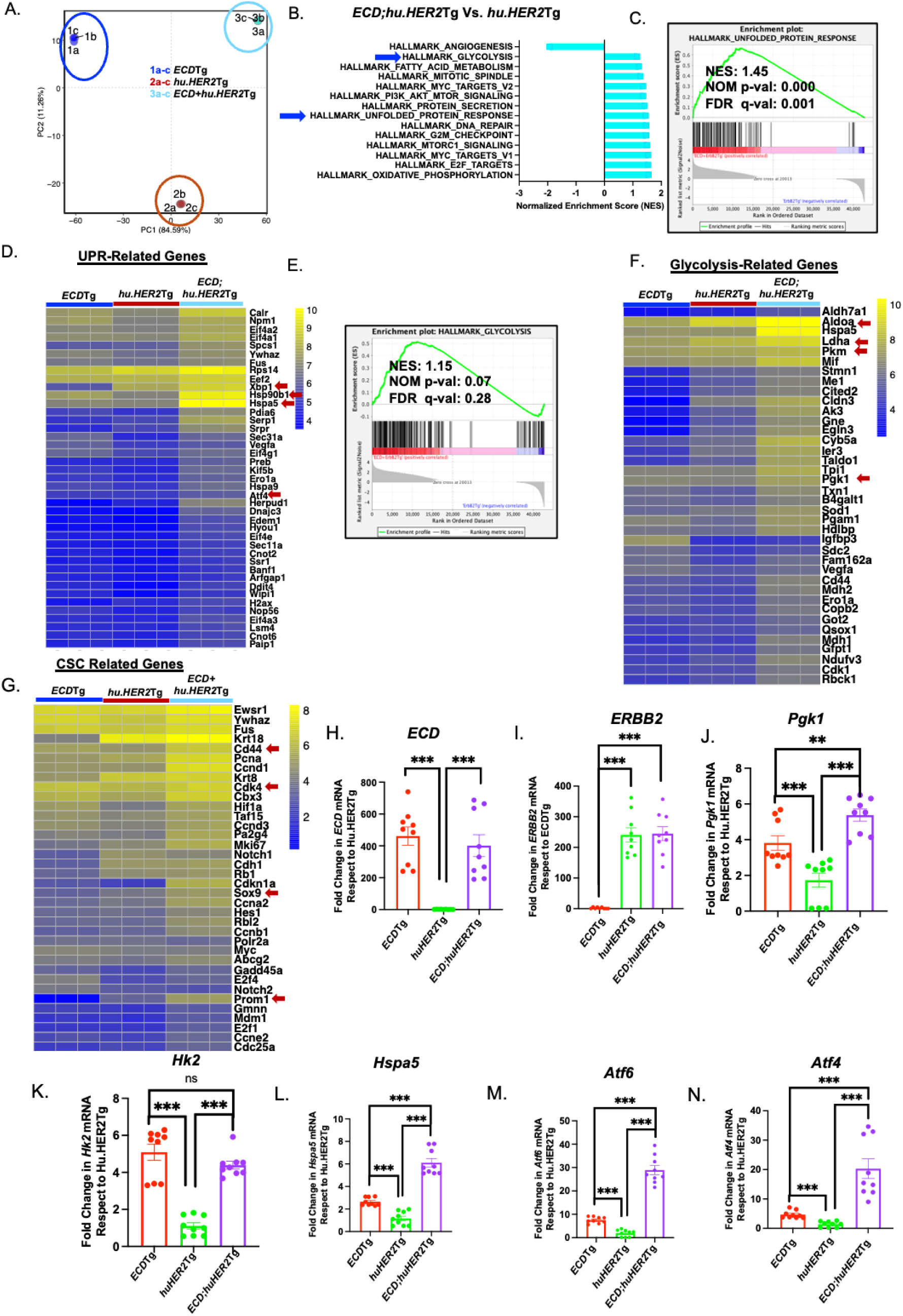
RNA-seq analyses of *ECD*Tg, *huHER2*Tg, and *ECD;huHER2*Tg tumors shows upregulation of two major oncogenic pathways, glycolysis and unfolded protein response in double transgenic tumors. **(A),** Principal component analysis (PCA) analysis of RNA-seq data shows clustering of *ECD*Tg tumors, *huHER2*Tg tumors and *ECD;huHER2*Tg tumors datasets. **(B)** Bar plot displays enriched hallmark gene sets. X-axis represents normalized enrichment scores (NES) of the signaling pathways with significant nominal p-values (NOM p-val). Blue arrows indicate unfolded protein response (UPR) and glycolysis (followed further in this study). **(C)** Enrichment plot of UPR signaling is presented with NES, NOM p-val and false discover rate q-value (FDR q-val). **(D)** Heatmap shows upregulated (yellow) and downregulated (blue) key UPR genes marked as red arrows. **(E)** Enrichment plot of glycolysis in *ECD;huHER2*Tg tumors compared to *huHER2*Tg tumors. NES, NOM p-val and FDR q-val. are indicated on the plot. **(F)** Heatmap of Key glycolytic genes is presented. Yellow represents upregulated and blue as downregulated genes. Major glycolytic genes are marked as red arrows. **(G)** Heatmap shows upregulation of key cancer stem cells related genes marked as red arrows in *ECD;huHER2*Tg tumors. **(H &I)** qRT-PCR confirms the presence of expected human *ECD and HER2* in tumors. **(J-N)** qRT-PCR analyses of indicated glycolytic and UPR genes. mRNA quantitation data represents mean ± SEM with two-tailed unpaired *t* test. *n* = 3; ns, *P* > 0.05; **, *P* < 0.01; ***, *P* < 0.001.

Further analysis of the RNA-seq data showed an upregulation of cancer stem cell (CSC) associated gene signature in DTg tumors as compared to single Tg tumors, including *CD44*, *PROM1*, *SOX9* and *CDK4* (**Fig. 3G and Supplementary Fig. S1J-M**). These findings are consistent with the key roles of cancer stem cells (CSCs), and EMT (43, 44) in mammary tumorigenesis and metastasis. RNA-seq analysis also showed the upregulation of EMT-related genes *CALD1, SAT1, PPIB, SNTB1*, *FERMT2 and MCM7* in DTg tumors (**Supplementary Fig. S1B)**. qRT-PCR analysis, in addition to confirming the expected overexpression of *ECD* and *HER2* in Tg tumors (**Fig. 3H &I**), validated the significant upregulation of glycolytic genes (*PGK1, HK2),* as well as UPR pathway related genes (*HSPA5*, *ATF6* and *ATF4)* in DTg tumors (**Fig. 3J-N**). Identification of the upregulation of UPR and glycolysis related gene expression in DTg tumors is consistent with the role of overexpressed ECD to mitigate ER stress (45) and to promote glycolysis-related gene expression to *ECD*Tg mice (19).

### Recapitulation of pro-oncogenic cooperation between ECD and ERBB2 in an immortal human mammary epithelial cell overexpression model

To experimentally demonstrate the co-oncogenic role of ECD and ERBB2 in human cells, we expressed the vector, ECD and/or ERBB2 in two distinct hTERT-immortalized human mammary epithelial cell lines (hMECs), 76NTERT and 70NTERT, which show no evidence of oncogenic transformation *in vitro* or *in vivo* in xenograft models (46). Western blotting demonstrated expected protein level of transfected ECD and ERBB2 (**Fig. 4A-F**).

**Fig. 4.**
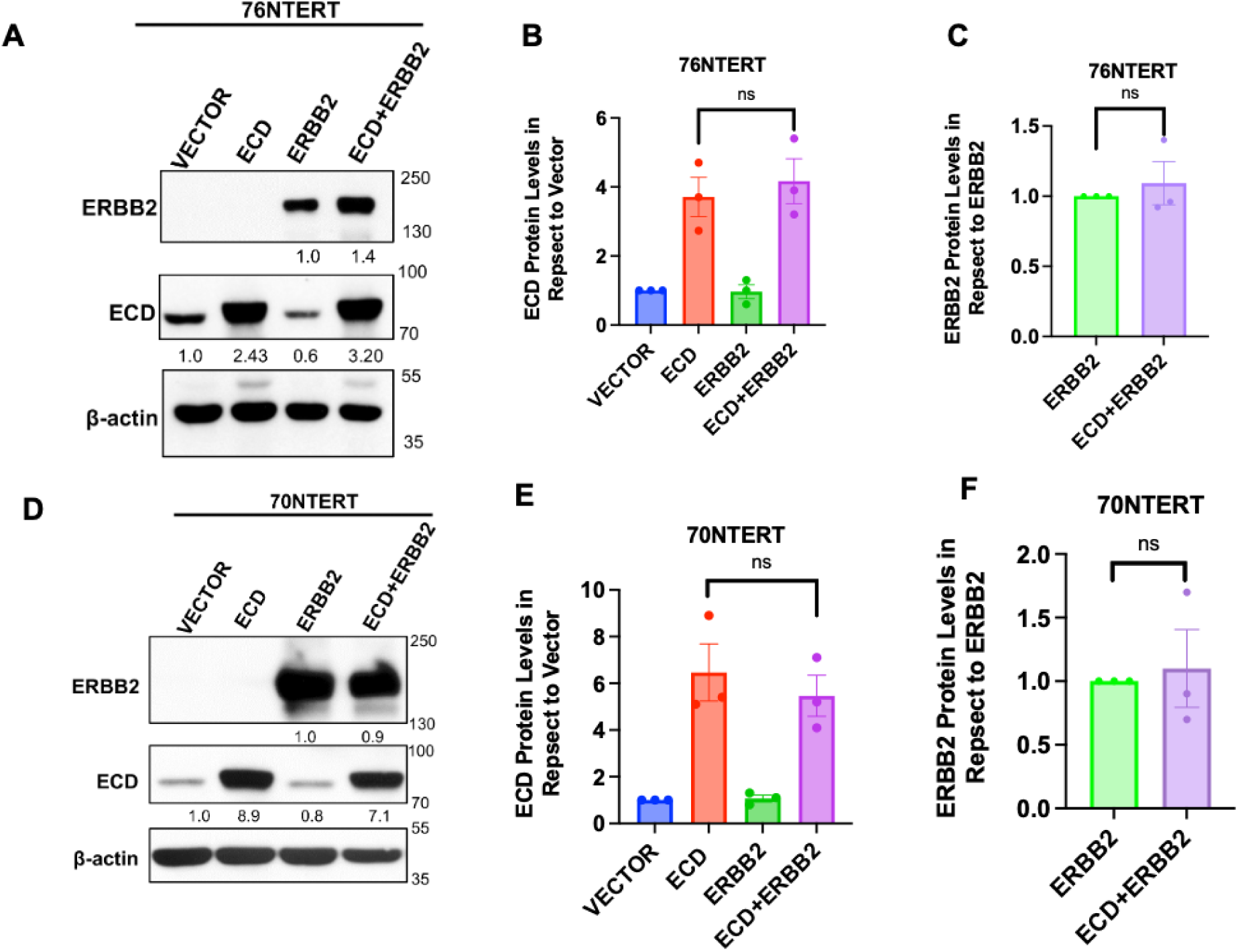
76NTERT and 70NTERT transductants stably overexpress ECD and ERBB2 protein and mRNA. **(A-F)** Western blotting of 76NTERT **(A-C)** and 70NTERT **(D-F)** expressing vector control, ECD, ERBB2, and ECD+ERBB2. Representative densitometries of ECD and ERBB2 are shown in respect to vector control and ERBB2 overexpressing cells respectively after normalization to loading control β-actin. Band intensities of three biological experiments were measured using ImageJ software and plotted. Data represents mean +/− SEM with two-tailed unpaired *t* test. *n* = 3; ns (not significant), Three technical replicates from three independent biological experiments.

Next, to examine the biological impact of ECD, ERBB2 or ECD+ERBB2 overexpression, we compared the *in vitro* oncogenic traits of the derived hMEC transductants. When cultured in starvation medium (DFCI-3) in a two-dimensional (2D) culture, the ECD overexpressing hMECs exhibited proliferation kinetics (measured using the cell-titer-Glo assay) comparable to that of the vector expressing controls, consistent with our prior findings (46). In contrast, ERBB2-overexpressing cells exhibit a significant increase in cell proliferation (**Supplementary Fig. S2A & B**), as has been reported with ERBB2 expressing MCF-10A, a spontaneously immortalized hMECs (47). Notably, the proliferation of ECD+ERBB2 overexpressing hMECs in 2D culture was comparable to that of ERBB2-overexpressing cells (**Supplementary Fig. S2A & B**). Notably, while ECD alone did not exhibit proliferative advantage in two-dimensional growth, a significant increase in anchorage-independent growth in soft agar was observed in both ECD or ERBB2-overexpressing hMECs **(Fig. 5A-C)**. Significantly, hMECs overexpressing ECD+ERBB2-exhibited a significantly more anchorage-independent growth in soft agar as compared to ECD− or ERBB2-alone overexpressing hMECs (**Fig. 5A-C**). Furthermore, mammosphere culture, a surrogate assay of stemness (28), showed similarly a significant increase in number and size of mammospheres in ECD+ERBB2-overexpressing cells vs. hMECs overexpressing only ECD or ERBB2 (**Fig. 5D-H**). Similar to soft agar colonies and mammospheres, ECD+ERBB2-overexpressing hMECs showed significantly larger sizes of organoids as compared to ECD− or ERBB2-overexpressing hMECs (**Fig. 5I-K)**. Finally, the ECD+ERBB2-overexpressing hMECs exhibited a significant increase in cell migration compared to ECD− or ERBB2-overexpressing hMECs (**Fig. 5L-N**). These results demonstrate that co-overexpression of ECD and ERBB2 promotes oncogenic traits in immortalized hMECs.

**Fig. 5.**
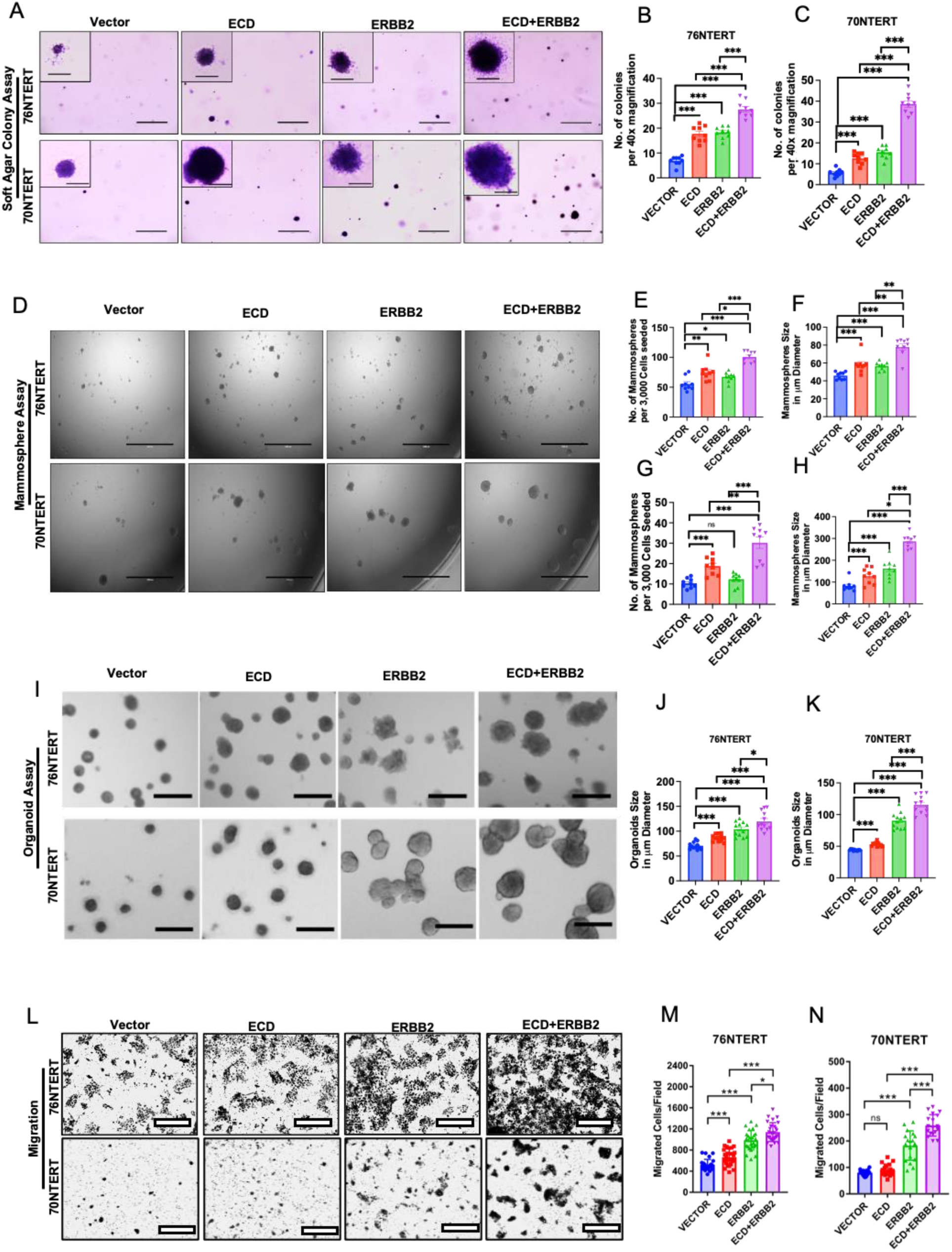
ECD and ERBB2 cooperate to promote anchorage independence, mammosphere formation, organoid formation and migration in immortal hMECs. **(A-C)** For anchorage independence assays six well plates were seeded with 20,000 cells per well in triplicates and stained with crystal violet after three weeks. Images show representative soft agar colonies for three biological experiments at 40x magnification (scale bar, 400 μm) with insets (magnification 200x, scale bar, 50 μm) 76NTERT (**upper panels**) and 70NTERT (**lower panels**) **(A).** Bar graphs show average number of soft agar colonies per field at 40x magnification (with a cut off size 20 μm) +/− SEM for 76NTERT (**B**) and 70NTERT (**C**) transductants in three biological experiments (***p<0.001). Statistics used unpaired *Student’s t-test* with Welch’s correction. **(D-F)** For mammosphere assays, a total of 3,000 cells plated per well of 96 well ultra-low attachment plates in mammosphere medium and imaged after day 5. Representative images of mammospheres in 76NTERT **(upper panels)** and 70NTERT (**lower panels**) transductants followed by quantitation of mammosphere numbers and sizes (76NTERT**, E &F;** 70NTERT**, G & H)**. Scale bar, 1000 µm (magnification 40x). **(I-K)** For organoid assays, a total of 5,000 cells plated per well in 4% Matrigel on top of 100% Matrigel layer in eight-well chamber slides for 5 days before imaging **(I)** Representative images of 3D organoid growth in Matrigel of 76NTERT (**upper panels**) and 70NTERT (**lower panels**) transductants on Day 5 followed by quantitation (76NTERT**, J) and (**70NTERT**, K**). Scale bar, 250 µm (magnification 200x). 50 μm diameter was size cut off of tumorspheres and organoids. Bar graph represents the size of organoids in µm diameter with mean ± SEM with unpaired *Student’s t-test* with Welch’s correction. *n* = 3 with four technical replicates; ^ns^p>0.05; *, *P* < 0.05; **, *P* < 0.01; ***, *P* < 0.001. **(L-N)** Transwell migration of cells. 20,000 cells per well were plated in DFCI-3 medium towards complete medium (DFCI-1) assayed after 18 hrs. (**L)** Representative images of 76NTERT (**upper panels**) and 70NTERT (**lower panels**) migrated cells (images at 100x magnification, white scale bar = 400 μm). Bar graph summarizes migratory capabilities of 76NTERT **(M)** and 70NTERT **(N)** transductants as average number of migrated cells +/− SEM per field at 100x magnification. Data represents unpaired *Student’s t-test* with Welch’s correction. *n* = 3 with three technical replicates; ^ns^ p>0.05; *p<0.05; **, *P* < 0.01; ***, *P* < 0.001

### RNA-Sequencing shows enrichment of Glycolysis and Unfolded Protein Response (UPR) Pathways in ECD+ERBB2 overexpressing hMECs

To assess if the transcriptomic alterations seen upon ECD and/or ERBB2 DTg could be recapitulated in the hMEC model, we performed RNA-sequencing comparison of vector, ECD, ERBB2 or ECD+ERBB2 overexpressing 76NTERT cells, each in triplicates. The principal component analysis (PCA) plot shows the expected non-overlapping clusters of the various cell lines analyzed (**Fig. 6A**). Notably, like analysis of the Tg model-derived tumors **(Fig. 3)**, the Gene Set Enrichment Analysis (GSEA) identified significant upregulation of the UPR, glycolysis, mTORC1 signaling, hypoxia signaling, and protein secretion signaling pathways in ECD+ERBB2 overexpressing hMECs in comparison with those overexpressing ECD or ERBB2 alone (**Fig. 6B-D**). The overexpression of *ECD* and *ERBB2* was confirmed from the transcript per million (TPM) values of the respective transductants (**Supplementary Fig. S3A & B**). In addition, we observed that hallmark E2F targets, glycolysis, mTORC1 signaling, hypoxia and EMT signaling were significantly enriched in ECD overexpressing cells as compared to vector cells (**Supplementary Fig. S3C**), and hallmark E2F targets, mTORC1 signaling and hypoxia pathways in ERBB2 overexpressing cells (**Supplementary Fig. S3D**). Notably, ECD+ERBB2 overexpressing hMECs showed higher expression levels of the EMT signature genes (*VIM, FOXA2, CD44, PPIB, FERMT2, TWIST2*) compared to single gene overexpressing cells (**Supplementary Fig. S3E, F and G)**. Further, the expression levels of several key genes related to the UPR pathway (*HSP90B1* (GRP94)*, ATF6, EIF2AK3* (PERK)*, ATF4*) (**Fig. 6E and Supplementary Fig. S3H-K**) and the glycolysis pathway *(LDHA, PGK1, HK2, SOD1, BIK, IDH1*) were higher in the ECD+ERBB2-overexpressing cells (**Fig. 6F and Supplementary Fig. S3L-Q**). qRT-PCR analyses confirmed the significant upregulation of glycolytic gene mRNAs, *HK2, LDHA, SLC2A1* (coding for GLUT1)*, ENO1* and *PDK1* in ECD− or ERBB2-overexpressing immortal hMECs and further upregulation in ECD+ERBB2 co-overexpressing cells (**Fig. 6G-K and Supplementary Fig. S4**). The qRT-PCR also confirmed the upregulation of UPR related genes, *HSPA5* (GRP78)*, ATF6* and *EIF2AK3* (PERK), in ECD+ERBB2 overexpressing vs. single gene overexpressing immortal hMECs (**Fig. 6L-N**).

**Fig. 6:**
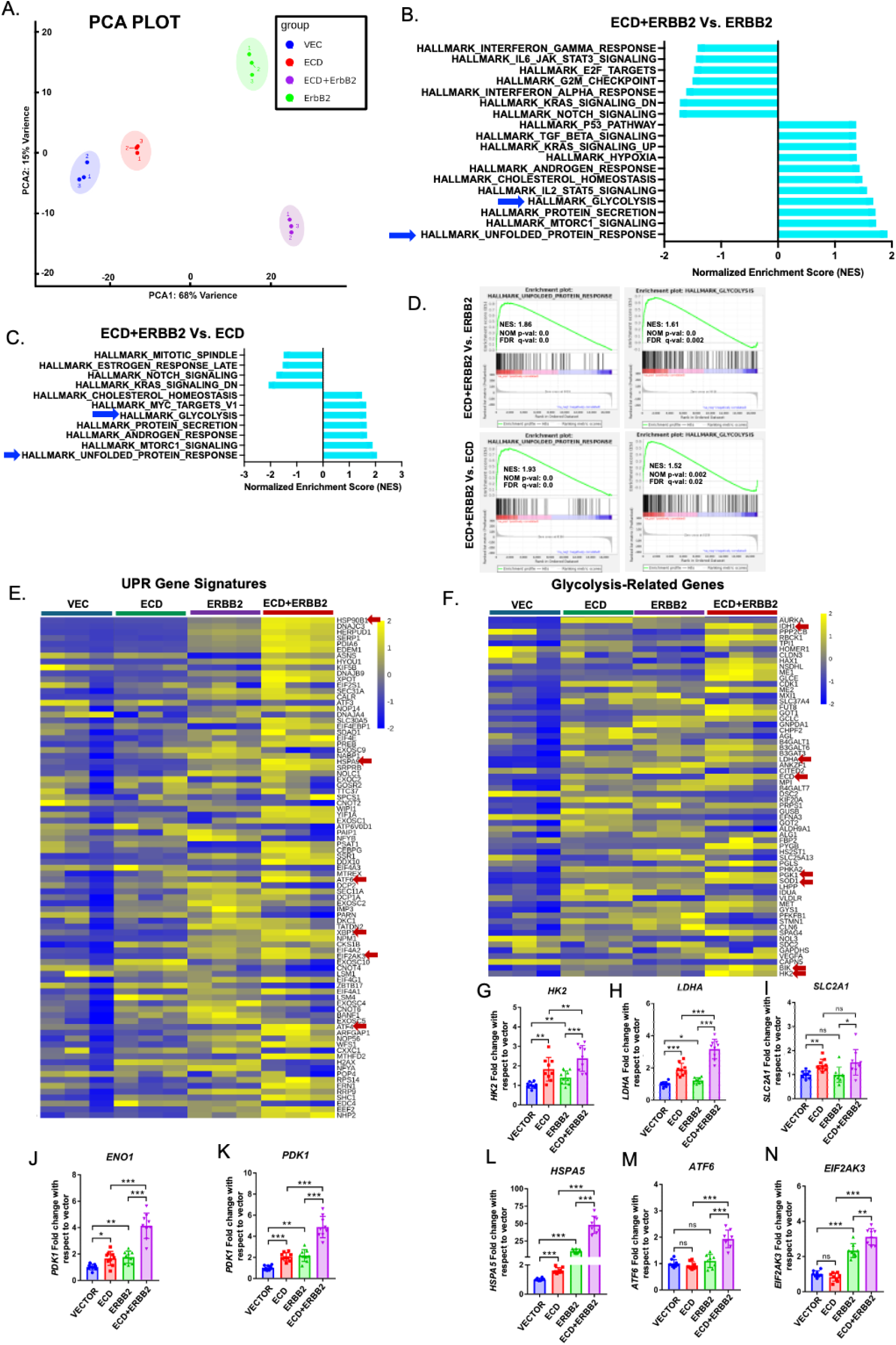
RNA-seq analyses of 76NTERT cells expressing vector, ECD, ERBB2 and ECD+ERBB2 shows upregulation of two major oncogenic pathways, glycolysis and unfolded protein response in ECD+ERBB2 cells. **(A)** Principal component analysis (PCA) analysis of RNA-seq data shows clustering of vector, ECD, ERBB2, and ECD+ERBB2 overexpressing 76NTERT cells. **(B, C)** Bar plots display enriched hallmark gene sets including unfolded protein response (UPR), and glycolysis. X-axis represents normalized enrichment scores (NES) of the signaling pathways with significant nominal p-values (NOM p-val). Blue arrows represent pathways emphasized in this study. **(D)** GSEA (Gene set enrichment analysis) plots display upregulation of UPR and glycolysis pathways genes in ECD+ERBB2 cells, as compared to ERBB2 or ECD overexpressing cells. NES, nominal p-values and false discovery rate (FDR) are indicated on the plot. **(E)** Heatmap shows upregulated (yellow) and downregulated (blue) key UPR genes (**E**) and glycolytic genes (**F**). qRT-PCR analyses of glycolytic genes *HK2* (**G**), *LDHA* (**H**), *SLC2A1* (GLUT1) (**I**), *ENO1* (**J**) and *PDK1* **(K)** in 76NTERT transductants are shown. mRNA quantitation data represents mean ± SEM with two-tailed unpaired *t* test. *n* = 3; ns, *P* > 0.05; **, *P* < 0.01; ***, *P* < 0.001. Relative UPR related mRNA levels of *HSPA5 (*GRP78) (**L**), *ATF6* (**M**), *EIF2AK3* (PERK) (**N**) are shown in indicated cells. Relative levels of each transcript are expressed as fold change with respect to vector control cells after normalizing with the house keeping gene, β-actin using the ΔΔCT method. Each bar graph indicates mean fold change +/− SEM from three experiments, each with three technical replicates (*p<0.05, **p<0.01, ***p<0.001).

### Functional validation of the UPR and glycolytic pathway upregulation in ECD+ERBB2 overexpressing hMECs

The UPR is a well-established pathway to play a role in cancer progression, maintenance, therapy resistance and metabolic rewiring (49). We have previously shown that ECD mitigates the negative consequences of ER stress by upregulating GRP78 levels (45). Overexpressed ERBB2 is also known to promote the activation of UPR (50).

Thus, we assessed the levels of GRP78 and sliced XBP1 (XBP1-s) in vector, ECD, ERBB2 or ECD+ERBB2 overexpressing 76NERT hMEC cell line, following growth factor deprivation to induce ER stress (51). Notably, the basal levels of GRP78 and XBP1-s were upregulated in ECD− (1.9-fold for GRP78 and 3.5-fold for XBP-1s) as well as ERBB2 (2.8-fold for GRP78 and 3.7-fold for XBP-1s) overexpressing 76NTERT cells in comparison to vector expressing cells. Significantly, the ECD+ERBB2 expressing cells showed further higher levels of the two proteins (3-fold for GRP78, 4.3-fold for XBP-1s). Notably, while GRP78 and XBP1-s levels were reduced within 24h of growth factor deprivation in vector or single gene transductants, their levels were maintained for substantially longer times in ECD+ERBB2 overexpressing 76NTERT cells (**Fig. 7A & B**). These results suggest that ECD+ERBB2 co-overexpression sustains the survival pathway by upregulating GRP78 and XBP-1s to maintain oncogenic traits.

**Fig. 7.**
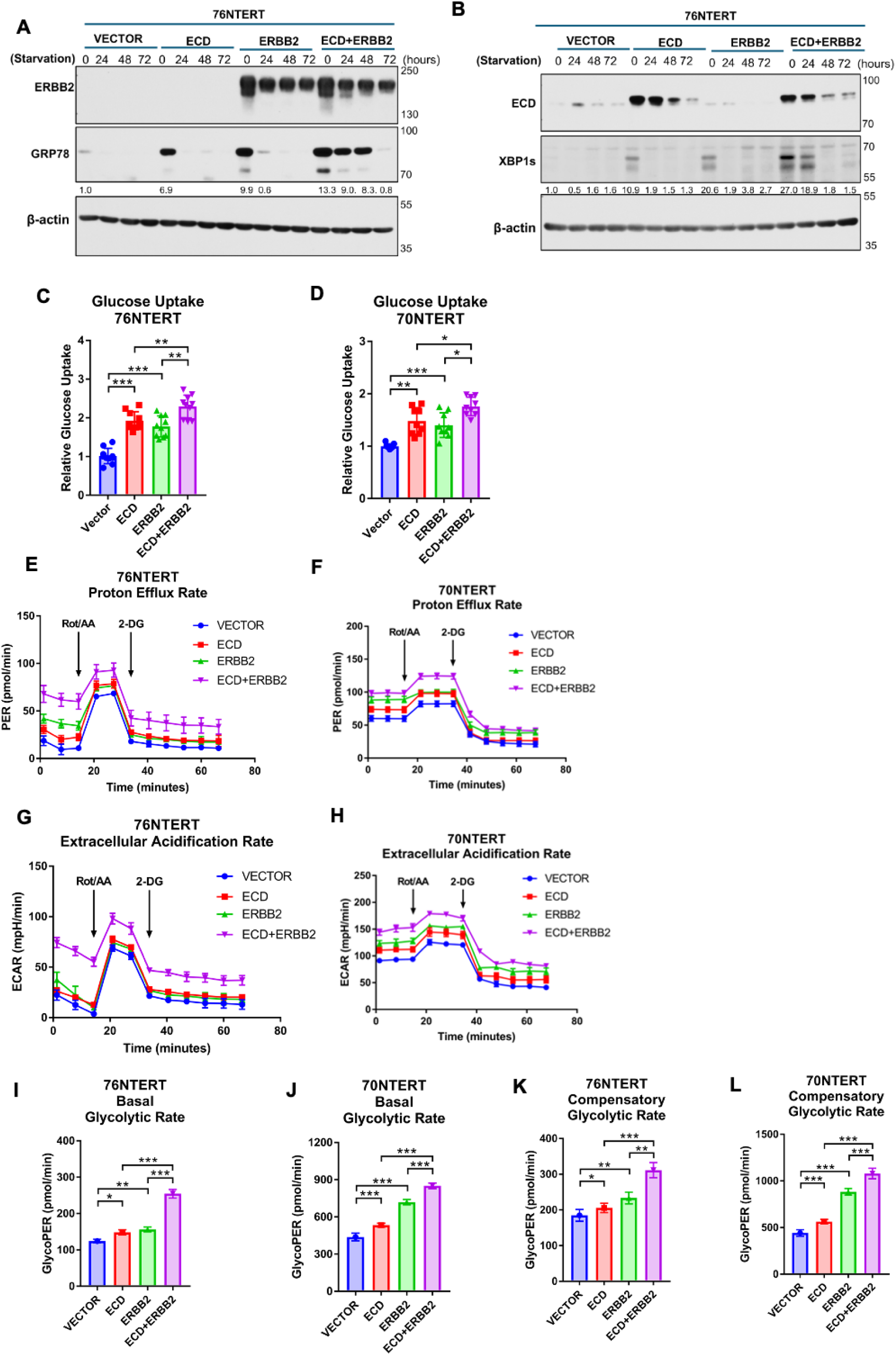
ECD and ERBB2 combined overexpression further enhanced UPR responses, glucose uptake, and glycolytic rate. **(A, B)** Western blotting of 76NTERT transductants grown in serum-free DFCI-3 medium at indicated time points revealed prolonged expression of GRP78 and XBP-1s (spliced form of XBP-1). ECD and ERBB2 expression confirms the overexpression of ECD and ERBB2 in transductants. Densitometries are in respect to no starvation of vector cells after normalizing with β-actin. Glucose uptake in 76NTERT **(C)** and 70NTERT **(D)** transductants. The values were normalized with respective to cell counts and depicted as compared with vector control. Quantification of results from three biological experiments, each with four technical replicates is shown as a bar graph. Data represents as mean ± SEM and two-tailed unpaired test with Welch correction (*p<0.05, **p<0.01, ***p<0.001). **(E-L)** Seahorse glycolytic rate was assessed in 76NTERT **(E)** and 70NTERT **(F)** transductants. Cells were seeded in 96-well plates and exposed to Rot/AA (rotenone and antimycin cocktail) and 2-DG to measure proton eflux rate **(PER, E & F)** and extracellular acidification rate **(ECAR, G & H)**. One representative PER and ECAR plots from three independent experiments are displayed (**E-H**). Basal glycolysis **(I & J)** and compensatory glycolysis **(K & L)** glycoPER in pmol/min are presented as bar graphs calculated from subtracting ECAR (extra cellular acidification rate) from PER (proton efflux rates) (**I-L**). Quantitation data represents mean ± SEM with two-tailed unpaired *t* test. *n* = 3; ns, *P* > 0.05; *, *P* < 0.05; **, *P* < 0.01; ***, *P* < 0.001.

To functionally link the upregulation of mRNA levels of glycolytic enzymes upon ECD, ERBB2 and ECD+ERBB2 overexpression, we assessed the glucose uptake in the hMEC models using the Promega Glucose Uptake-Glo method. A significant increase in glucose uptake was observed in ECD-overexpressing hMECs (**Fig. 7C & D**), consistent with our previous findings (19). As expected, ERBB2 overexpressing hMECs also exhibited an increase in glucose uptake, consistent with the literature (52). Significantly, hMECs co-overexpressing ECD and ERBB2 exhibited a further increase in glucose uptake (**Fig. 7C & D**), again emphasizing the co-operation of ECD and ERBB2 to maintain metabolism.

To assess a direct functional impact on glycolysis, we employed the Seahorse XF glycolytic rate assay to determine the proton efflux rate (PER) from glycolysis (GlycoPER) and extracellular acidification rate (ECAR) as readouts of glycolysis. Both the PER (**Fig. 7E & F**) and ECAR (**Fig. 7G &H**) measured in the presence of rotenone (Rot) and antimycin A (AA) to inhibit mitochondrial respiration, were increased in immortal hMECs overexpressing ECD or ERBB2 and more so in cell overexpressing ECD+ERBB2. Subsequent inhibition of hexokinase with 2-DG confirmed the measured parameters are due to glycolysis. Notably, the ECD, ERBB2 or ECD+ERBB2 overexpressing immortal hMECs exhibited significantly higher levels of basal glycolysis (GlycoPER data for first 15 mins prior to Rot/AA addition) and increased glycolytic capacity (difference between pre- and post-Rot/AA levels of PER), glycolytic reserve (difference between glycolytic capacity and glycolysis rate), and compensatory glycolysis (GlycoPER data after 2-DG addition) (**Fig. 7I-L**). These findings support a cooperative role of ECD and ERBB2 to promote the glycolytic pathway in hMECs.

Our recent findings identified ECD as an RNA binding protein (21). The enhanced-CLIP identified ECD binding to target mRNAs including the mRNAs of glycolytic gene *PKM, a* UPR related gene *HSPA5*. RNA immunoprecipitation (RIP) assay to assess if ECD associated with mRNAs confirmed that ECD forms a complex with the mRNAs of key glycolytic enzymes (*HK2, PKM*, *LDHA)* and UPR-associated genes (*HSPA5,* GRP78 gene name) (**Fig. 8A)**. In view of the increased levels of glycolysis and UPR-associated gene mRNAs upon ECD+ERBB2 overexpression, and the association of ECD with these mRNAs, we examined the stability of *HK2*, *LDHA, PKM*, and *HSPA5* transcripts in various hMECs transductants by assessing the half-life of mRNAs after inhibiting global transcription with actinomycin D. Notably, in comparison to vector-transduced hMECs, a significant increase in the half-life was observed in hMECs overexpressing ECD [fold increase-HK2 (1.6); LDHA (1.5); PKM2 (2.0)] or ERBB2 [fold increase-HK2 (1.7); LDHA (1.7); PKM2 (1.6)], and the increase was significantly more pronounced in ECD+ERBB2 hMECs transductants [HK2 (3.1); LDHA (2.8); PKM2 (3.0)] (**Fig. 8B-G**). While the *HSPA5* mRNA stability was unchanged by the overexpression of ERBB2 alone as compared to vector-expressing immoral hMECs (**Fig. 8H & I**), a significant increase was observed in immortal hMECs overexpressing ECD (fold increase 2.0) or ECD+ERBB2 (fold increase 2.4) (**Fig. 8H & I**). These results indicate that overexpression of either ECD or ERBB2 increases the mRNA stability of glycolytic enzymes, and co-overexpression of ECD+ERBB2 produces an additive effect. Moreover, ERBB2 expression alone did not affect the stability of HSPA5 mRNA, while ECD overexpression did promote the mRNA stability, suggesting that the combined effect of ECD+ERBB2 overexpression on GRP78 protein levels (**Fig. 8H & I**) may involve additional effects on protein stability, in addition to mRNA stabilization contributed by ECD.

**Fig. 8.**
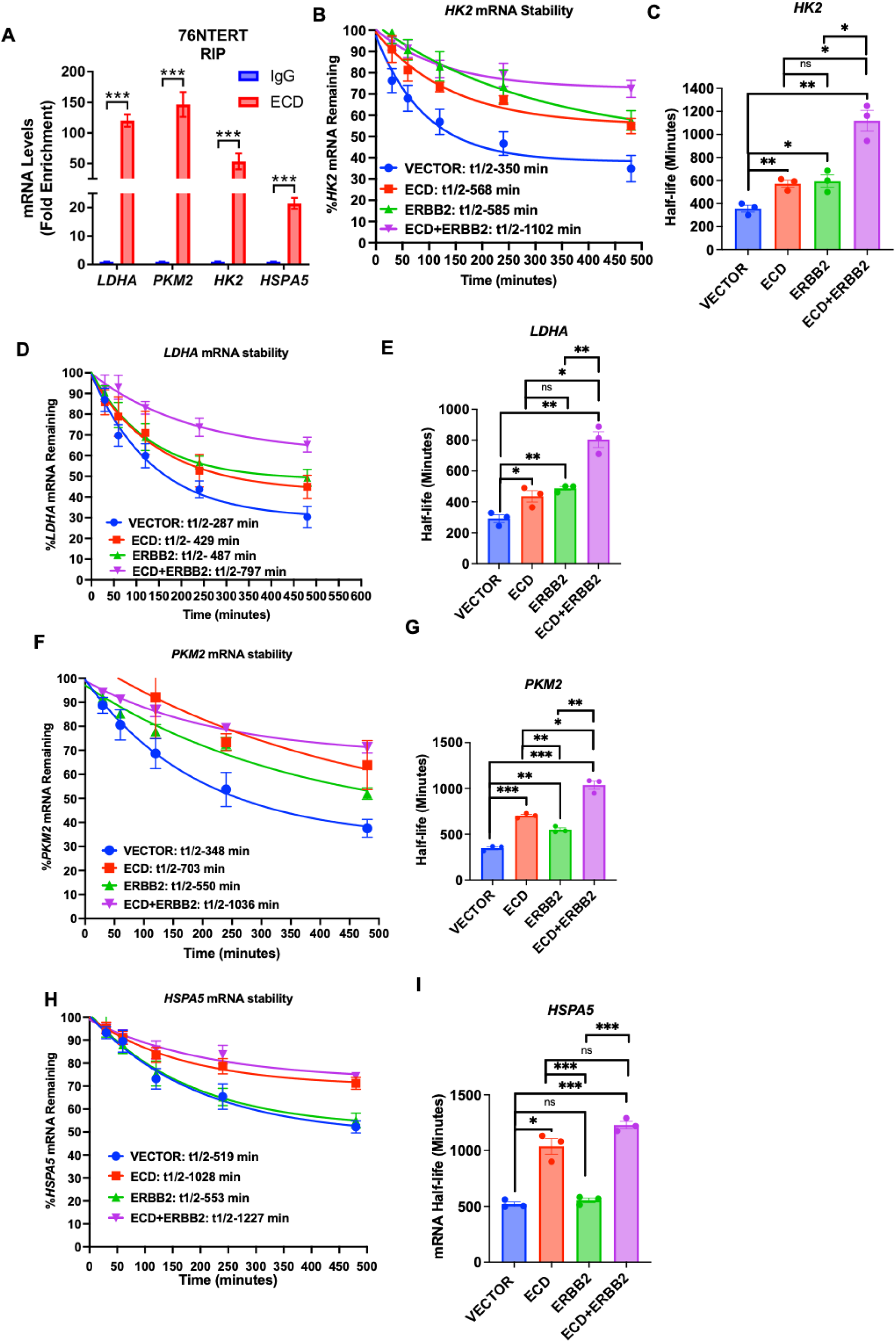
ECD protein associates with indicated mRNAs and ECD overexpression promotes stability. **(A)** ECD association with glycolytic and UPR-related mRNAs are assessed by RNA immunoprecipitation assay. Cell lysates were collected in immunoprecipitation lysis buffer. RNA-protein complexes were isolated using anti-ECD and anti-IgG antibodies. RNA was isolated and reverse-transcribed to make cDNA. Relative levels of indicated mRNAs were measured by qRT-PCR. Graph displayed mean fold enrichment in each respective gene, as compared to IgG control. Quantitation data represents mean ± SEM with two-tailed unpaired *t* test. *n* = 3; ***, *P* < 0.001. **(B-I)** 76NTERT transfectants were treated with actinomycin D for indicated time points to inhibit transcription. Trend lines were created by measuring relative abundance of each transcript with respect to the 0-minute time point for each transfectant and are expressed as percentage of mRNA remaining. Transcript abundance was measured by qRT-PCR using 18S rRNA as a housekeeping gene. Half-life (t1/2) for indicated mRNAs are displayed for each transfectant. Data represents as mean ± SEM and two-tailed unpaired test with Welch correction (n=3 with three technical replicates; ns, *p* > 0.05; *p<0.05, **p<0.01, ***p<0.001).

## DISCUSSION

While prior studies demonstrating the overexpression of ECD in ERBB2-positive BC patients and the positive correlation between ECD overexpression and shorter patient survival (17, 18) strongly supported an important role for ECD in promoting ERBB2-driven oncogenesis, the nature of the ECD-ERBB2 cooperation and its mechanism were unknown. In this study, we established *in vivo* double *ECD;huHER2* transgenic mice model, and *in vitro* hTERT immortal hMEC expressing single or double genes models to elucidate the pro-oncogenic function of ECD with ERBB2. Our studies provide direct evidence that co-overexpression of ECD with ERBB2 promotes oncogenesis. Mechanistically, RNA-seq analysis and ECD-mRNA interaction analyses demonstrate that the ECD binds to and promotes the stability of mRNAs for key components of glycolysis and UPR to upregulate their levels, thereby promoting the activity of these critical pro-oncogenic cellular pathways in ECD+ERBB2 expressing mammary cells.

We began our study by crossing the previously published *ECD*Tg (19) mice with *MMTV-huHER2*Tg mice (20), we generated a unique double transgenic mouse model in which the overexpression of both human ECD and human ERBB2/HER2 (huHER2) is targeted to the mammary epithelium. Notably, co-overexpression of ECD did not alter the latency (7-11 months) or incidence (38% in *ECD;huHER2*Tg tumors vs 32% in *huHER2*Tg) of tumors compared to *huHER2*Tg mice, however the *ECD;huHER2*Tg mice developed heterogenous mammary tumors exhibiting aggressive features of adenosquamous carcinoma, and spindle cell carcinoma with EMT features in addition to solid nodular carcinoma (**Supplementary Table S3**), consistent with heterogenous tumors seen in HER2+ breast cancer patients (53, 54). Our observation of *huHER2*Tg mice producing solid nodular carcinoma (**Fig. 1)** is consistent with a published report (38). Notably, the heterogenous features of *ECD;huHER2*Tg tumors resemble the features we observed in *ECD*Tg mice (19), supporting the likelihood that the altered features of *ECD;huHER2*Tg tumors are due to ECD overexpression. Furthermore, the *ECD;huHER2*Tg tumors exhibited a higher proliferative index, a feature of aggressive tumors, compared to *huHER2*Tg tumors (**Fig. 2**). Notably, the *ECD;huHER2*Tg tumors expressed high cytokeratin marker 14 (CK14), nuclear p63 and luminal cytokeratin marker, CK18 suggesting intra-tumoral heterogeneity (**Fig. 2**) and expression of basal mammary epithelial markers, indicating a switch towards basal-like tumor phenotype, resembling human ERBB2 tumors with intra and inter tumor heterogeneity (54).

Increasing evidence suggest that tumor heterogeneity and aggressiveness reflects the functional impact of cancer stem cells (CSCs) in tumors (43, 55). Prior studies have shown a key role of CSCs in ERBB2-driven tumorigenesis (56). Interestingly, we have found that ECD overexpression promotes mammosphere growth (19), a surrogate assay of BC CSCs (43, 55). Our transcriptome profiling analyses of *ECD;huHER2*Tg tumors revealed an enrichment of CSC-related gene expression in comparison with *huHER2*Tg or *ECD*Tg tumors (**Fig. 3**). Both *ECD*Tg (19) and *huHER2*Tg (41) tumors show increased expression of EMT markers, and EMT itself is linked to the CSC phenotype in tumors (44). Consistent with this idea, *ECD;huHER2*Tg tumors exhibited strong expression of EMT marker proteins using both immunohistochemistry and transcriptome analyses (**Fig. 2 and 3**). The oncogenic co-operation between ECD and ERBB2 was recapitulated in an *in vitro* model where we overexpressed ECD and/or ERBB2 in TERT-immortalized hMECs 76NTERT and 70NTERT (**Fig. 4**), which by themselves exhibit a phenotype akin to normal mammary epithelial cells (23, 46). We found that co-overexpression of ECD and ERBB2 led to enhanced migration, mammosphere, and organoid growth as well as anchorage independent growth in soft agar (**Fig. 5**). Interestingly, overexpression of ECD in hMECs did not provide a proliferative advantage in 2D cultured hMECs grown in growth factor-deprived (DFCI-3) medium nor did its co-overexpression with ERBB2 further enhance the higher cell proliferation/survival imparted by ERBB2 alone overexpression (**Supplemental Figure S2**). The lack of an impact of ECD overexpression in 2D hMEC culture on plastic (**Supplemental Figure S2**) vs. an enhancement of growth in 3D contests (**Fig. 5**) is consistent with higher ki67 staining seen in *ECD; huHER2*Tg tumors (**Fig. 2**). This discordance suggests that ECD may impinge on mechanisms critical for tumor growth in a natural 3D environment. In this regard, the ECD-regulated processes of glycolysis and UPR (19, 57, 58) are known to provide pro-survival mechanisms encountered in the harsh tumor microenvironments modeled *in vitro* in 3D. Overall, the pro-oncogenic behavior of ECD+ERBB2 vs. ERBB2-overexpressing hMECs is consistent with our findings in the Tg models. Together, the *in vivo* and *in vitro* oncogenesis studies support the role of ECD overexpression as a pro-tumorigenic adaptation in ERBB2-driven cancer as suggested by correlative studies of ECD overexpression in ERBB2+ patients (17, 18).

Recapitulating the results of in *ECD;huHER2*Tg tumors (**Fig. 3**), in the RNA sequencing analysis of 76NTERT hMEC cell models showed an upregulation of glycolysis and UPR associated gene expression in ECD+ERBB2 co-overexpressing hMECs vs. ERBB2 or ECD overexpressing cells (**Fig. 3 & 6**). Direct measurements demonstrated that overexpression of ECD with ERBB2 indeed promotes a high rate of glycolysis, which is accompanied by increased uptake of the requisite fuel glucose (**Fig. 7**). The latter is consistent with an increase in mRNA expression for GLUT1 (*SLC2A1*) (**Fig. 6**). Our analyses in the hMEC models also validated the increased expression of UPR-associated GRP78 (*HSPA5*) and the hMECs co-overexpressing ECD and ERBB2 showed a more sustained GRP78 expression under growth factor deprivation stress (**Fig. 7A & B**).

Reprogramming of glucose metabolism is an important hallmark of cancer and supports tumor cell growth and survival as well as remodeling of tumor microenvironment during cancer progression and metastasis (59). ECD-dependent enhancement of glycolysis in the context of ERBB2-driven oncogenesis in the present study is consistent with *ECD*Tg-derived tumor organoid models and cell-based analyses demonstrating increased glucose uptake upon ECD overexpression (19). ERBB2-driven oncogenesis itself is known to rewire glycolytic metabolism. Previous studies have shown that ERBB2 overexpression promotes breast cancer cell glycolysis and growth through heat shock factor 1-dependent increase in LDHA mRNA expression (52). ERBB2 overexpression dependent constitutive activation of PI3K/AKT signaling is known to induce the expression of hypoxia-inducible transcription factor HIF1α (60), resulting in transcriptional upregulation of glycolytic enzymes including LDHA (52).

Transformation by oncogenes, including ERBB2, induces ER stress and the associated UPR (14), appears to be required for tumor cell survival (11) and has been linked to resistance to ERBB2 targeted therapies (61). Since ER stress and the associated UPR induce a block in protein translation, promote mRNA degradation and induce a cell cycle block or apoptosis, tumor cells with persistent ER stress and UPR must turn on adaptive pathways to mitigate the negative consequences of ER stress. We have previously established that ECD overexpression protects cells from apoptotic outcome of ER stress induction, and this is mediated by ECD-dependent upregulation of GRP78 levels (45). The upregulation of GRP78 and its persistence (and that of and the spliced form of XBP-1) during growth factor deprivation support a similar role of ECD to promote ER stress tolerance in the context of ERBB2-driven oncogenesis without impairing the pro-oncogenic drive of ERBB2.

Notable overexpression of ECD further increases the mRNA expression of glycolytic and UPR genes, both in Tg tumors and in the hMEC system (**Fig. 6** and **Supplementary Fig. S3 & S4**), suggests the possibility that ECD may contribute to this process through mechanisms complementary to those by which ERBB2 alters these processes. In this regard, we and others have shown that ECD plays a key role in promoting mRNA biogenesis through its interactions with components of the mRNA splicing and nuclear to cytoplasmic mRNA export (17). It is plausible that ECD promotes these processes to complement the ERBB2-driven transcriptional activation of glycolytic and UPR gene expression. Consistently, our recent identification of ECD as a novel RNA binding protein (21), suggested the possibility that ECD my interact with the mRNAs of glycolytic and UPR genes whose expression is upregulated because of ECD overexpression. Indeed, the corresponding mRNAs were included among the ECD-bound mRNAs identified by enhanced CLIP-seq analysis, and our mRNA co-immunoprecipitation (RIP) assays confirmed the association of ECD with the mRNAs of *HK2*, *LDHA*, *PKM* and *HSPA5* (**Fig. 8**). Consistent with our recent findings that association with ECD stabilizes its target mRNAs (17, 19), hMECs with ECD overexpression exhibit increased stability of the ECD target mRNAs involved in glycolysis and UPR (**Fig. 8**). The enhanced stability of ECD-bound mRNAs suggests a novel and plausible mechanism by which ECD overexpression enhances glycolysis and pro-survival outcomes of UPR during ERBB2 driven breast cancer. Thus, our findings are consistent with complementary mechanisms by which ECD (via mRNA stability) and ERBB2 (via increased transcription) upregulate the expression of key genes in glycolysis, UPR and other pathways to sustain the tumorigenic drive. These mechanisms likely work in concert with others previously defined ones such as ERBB2-driven phosphorylation of Tristetraprolin (TTP/ZFP36) to inhibit the AU-rich element-mediated mRNA decay to promote mRNA stability (62).

We note that GSEA pathway analysis of RNA-seq data in *ECD;huHER2*Tg tumors also showed upregulation of additional pathways of including cell proliferation-related gene sets (E2F targets, G2M checkpoint, MYC target v1 and v2, and MITOTIC spindle), as well as MTORC1 signaling and DNA repair. Future studies will explore the role of these pathways in oncogenic cooperation between ECD and ERBB2.

In conclusion, we demonstrate a novel role co-overexpression role of ECD with ERBB2 in promoting breast cancer oncogenic drive. Our studies implicate two key tumor progression pathways, upregulation of glycolysis and UPR, as important contributors to ECD+ERBB2 cooperation, and identify the direct ECD binding and stabilization of mRNAs coding for key genes in the implicated pathways as a novel mechanism for how ECD may contribute to ERBB2-driven tumorigenesis. Given the known involvement of glycolysis and UPR in promoting oncogenesis, our studies provide key mechanistic insights into how ECD overexpression may specify poor prognosis and shorter survival in ERBB2+ breast cancer patients (17, 18).

## ACKNOWLEDGEMENTS

This work was supported by the NIH grants R21CA241055 and R03CA253193 to V. Band, and R35GM147467 to MJ Rowley; Department of Defense grants W81XWH-17–1-0616 and W81XWH-20–1-0058 to H. Band, and W81XWH-20–1-0546 and HT94252410337 to V. Band; Nebraska Initiative Grants to V. Band and B. Mohapatra. Support for the University of Nebraska Medical Center Genomics (Omaha, NE) was from NCI Support Grant P30CA036727 to the Fred and Pamela Buffett Cancer Center (Omaha, NE).

## AUTHOR CONTRIBUTIONS

Conceptualization and design; B.B.K., M.R., B.C.M., H.B., and V.B.

manuscript writing; B.B.K., M.R., B.C.M., H.B., and V.B.

data analysis and interpretation. B.B.K., M.R., S.M., A.R.R., F.O., M. S.M.., S.M.L., T.E.R., L.L., M.J.R., S.W., B.C.M., H.B., and V.B.

manuscript editing; B.C.M., H.B., and V.B.

bioinformatics analysis for the manuscript; L.L., S.W., T.E.R., M.J.R.

All authors have read and approved the final version of the manuscript and agree with the order of presentation of the authors.

---

**Supplementary Table-S1:**
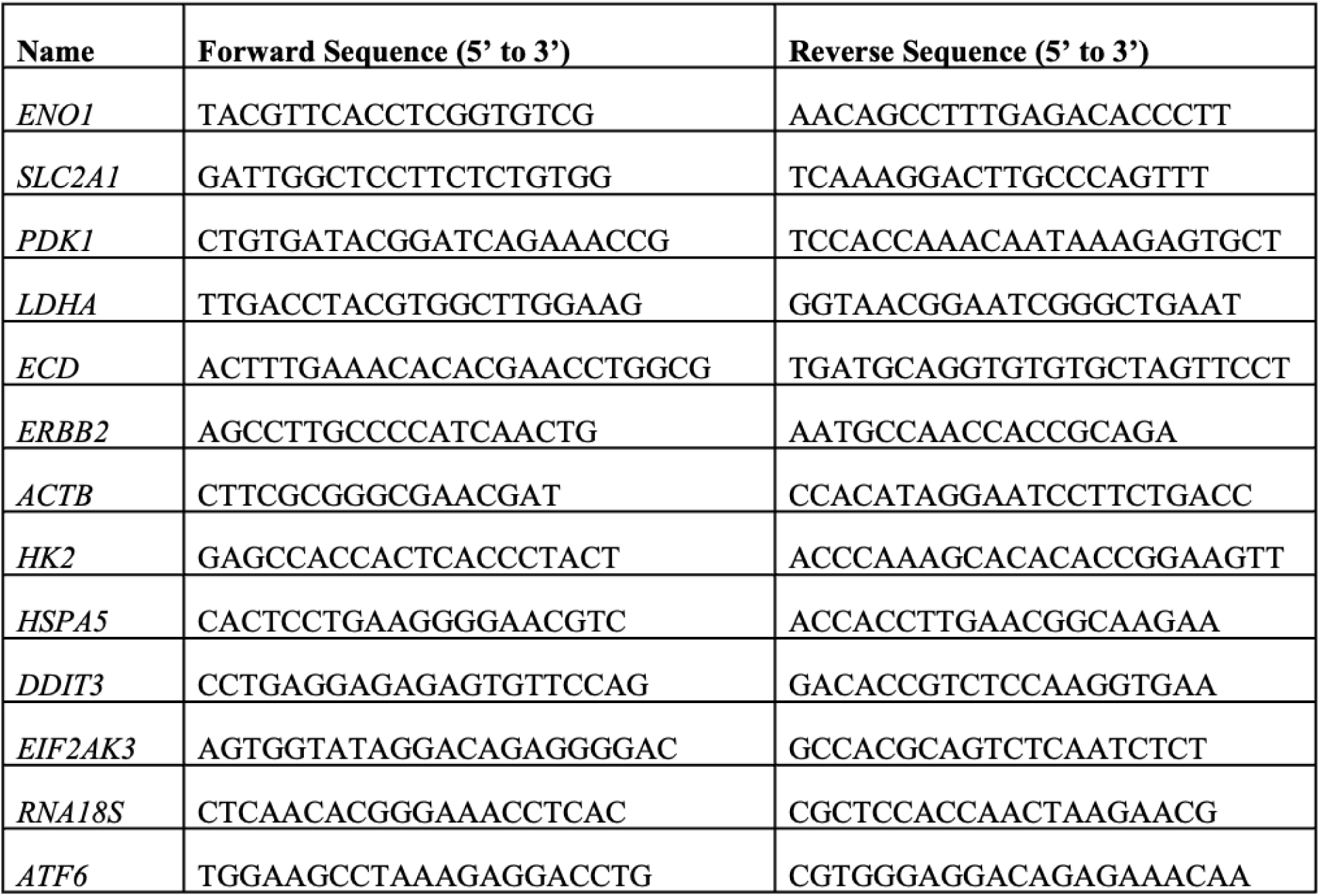
List of real time qPCR primers of human genes.

**Supplementary Table-S2:**
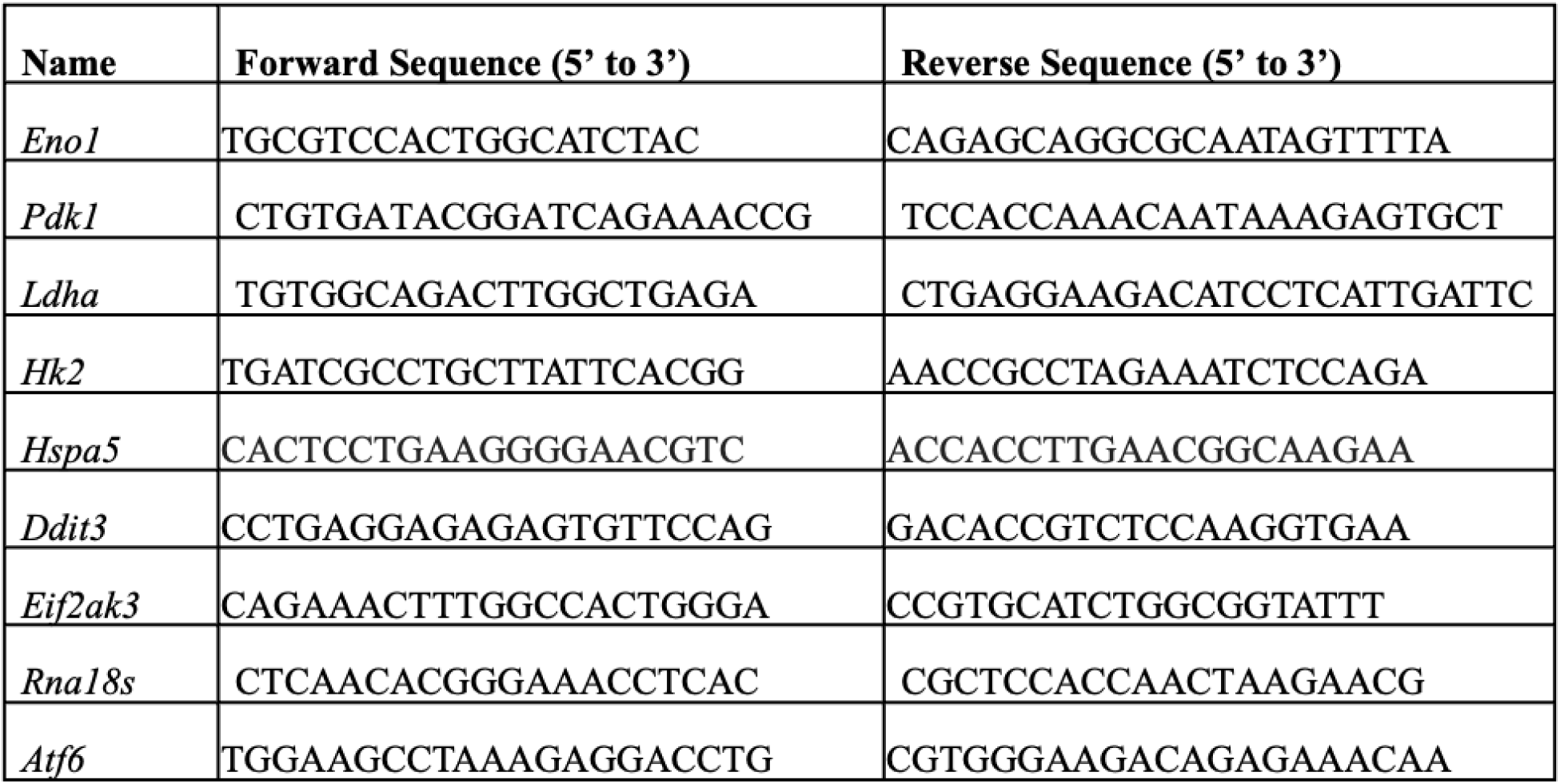
List of real time qPCR primers of mouse genes.

**Supplementary Table-S3:** Tumoe Latency and Incidence in MMTV.*huHER2* and *Tet(O)-ECD*; *MMTV-tTA*; *MMTV.huHER2* Transgenic Mice.

| **Genotype** | **Total**<br>**number of**<br>**mice** | **Number (%)**<br>**with mammary**<br>**tumors** | **Latency range**<br>**(Months)** |
| --- | --- | --- | --- |
| **MMTV.*huHER2*** | 34 | 11/34 (32%) | 7-11 |
| ***Tet(O)-ECD ;***<br>***MMTV-tTA*** | 51 | 17/51 (33%)* | 15-25* |
| ***Tet(O)-ECD ;***<br>***MMTV-tTA;***<br>***MMTV.huHER2*** | 21 | 8/21 (38%) | 7-10 |

(\* Data from Mohapatra/Mirza et al., MCR, 2022)

**Fig. S1:**
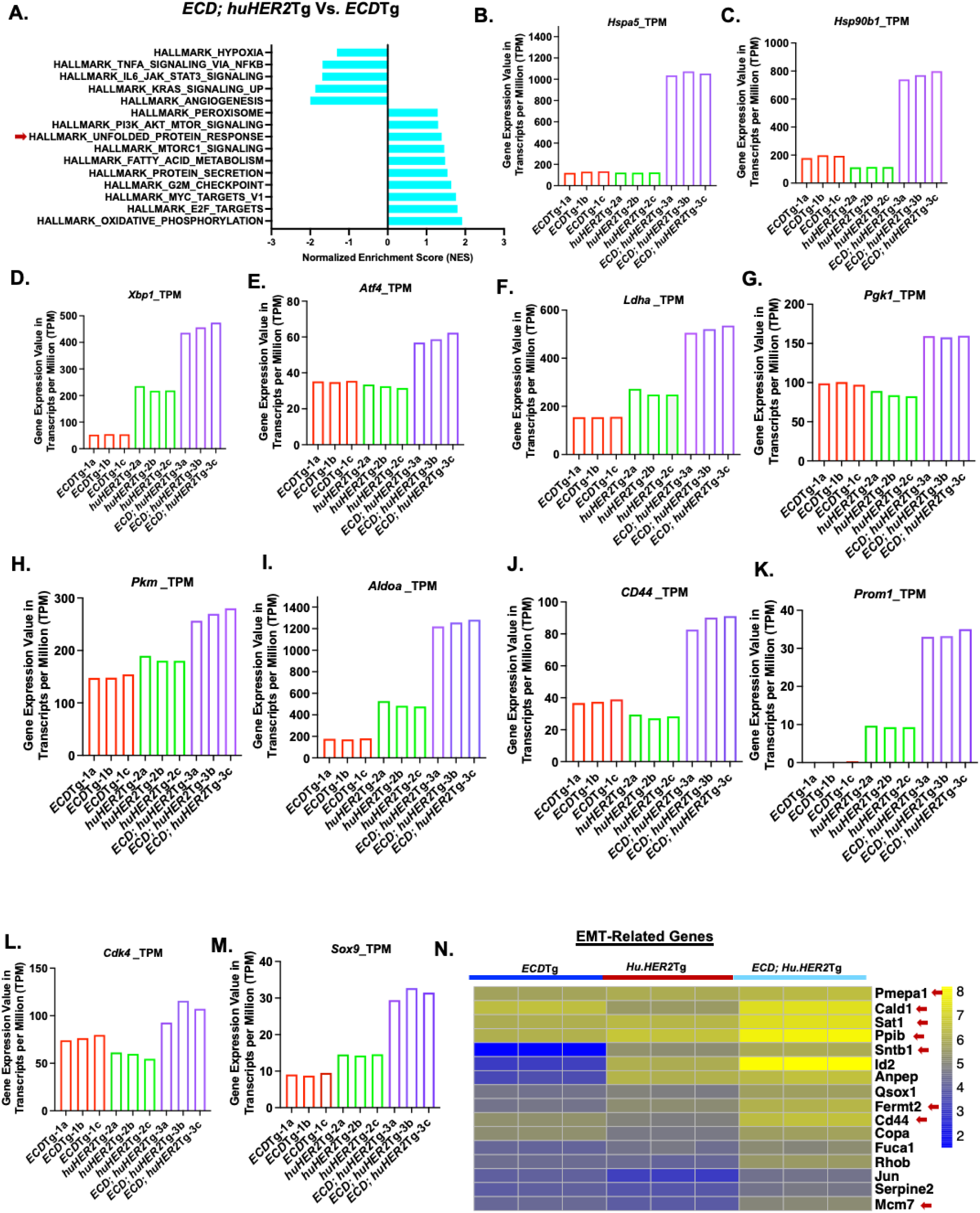
RNA-seq analyses of tumors from *ECD*Tg, *huHER2*Tg, and *ECD;huHER2*Tg mice display upregulation of UPR, glycolysis, CSCs and EMT-related genes: Bar plots display enriched hallmark gene sets including unfolded protein response (UPR), glycolysis, protein secretion, oxidative phosphorylation, mTORC1 signaling, E2F targets and PI3K-AKT-mTOR signaling in *ECD;huHER2*Tg tumors compared to *ECD*Tg tumors alone. X-axis represents normalized enrichment scores (NES) of the signaling pathways with significant nominal p-values (NOM p-val). Bar graphs represent gene expression values in transcripts per million (TPM) from the RNAseq analysis of all tumors. TPM values of UPR related genes *Hspa5* (B), *Hsp90b1* (C), *Xbp1* (D), *Atf4* (E), glycolysis related genes *Ldha* (F), *Pgk1* (G), *Pkm* (H), *Aldoa* (I) and CSCs related genes *CD44* (J), *Prom1* (K), *Cdk4* (L), *Sox9* (M) are displayed. (N) Heatmap shows upregulation of key EMT-related genes in *ECD;huHER2*Tg tumors.

**Fig. S2.**
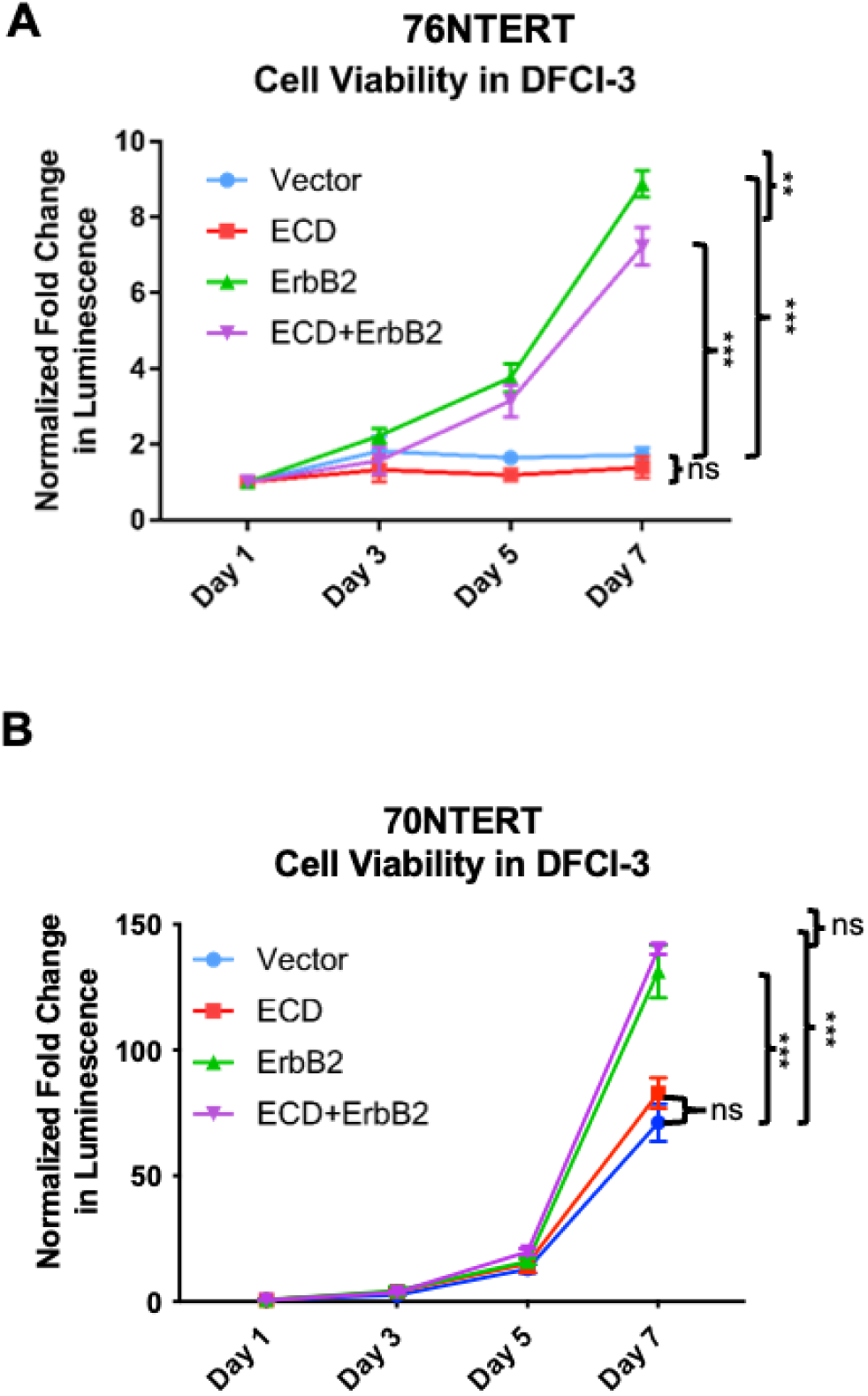
Cell viability over seven days of 76NTERT and 70NTERT transfectants as measured by CellTitre-Glo luminescence assay in starvation DFCI-3 medium (A & B). Cells seeded at a density of 1,000 cells per well in 96-well plates. Viability by luminescence readings taken at indicated time points over seven days. Luminescence values normalized to day 1 time point for each condition. Figure shows mean +/− SEM for three biological experiments each with 6 technical replicates per condition at each timepoint. Statistical significance between mean fold changes were calculated by two-way ANOVA (ns, p > 0.05; **, p < 0.01; ***, p < 0.001) in 76NTERT (A) and 70NTERT (B).

**Fig. S3.**
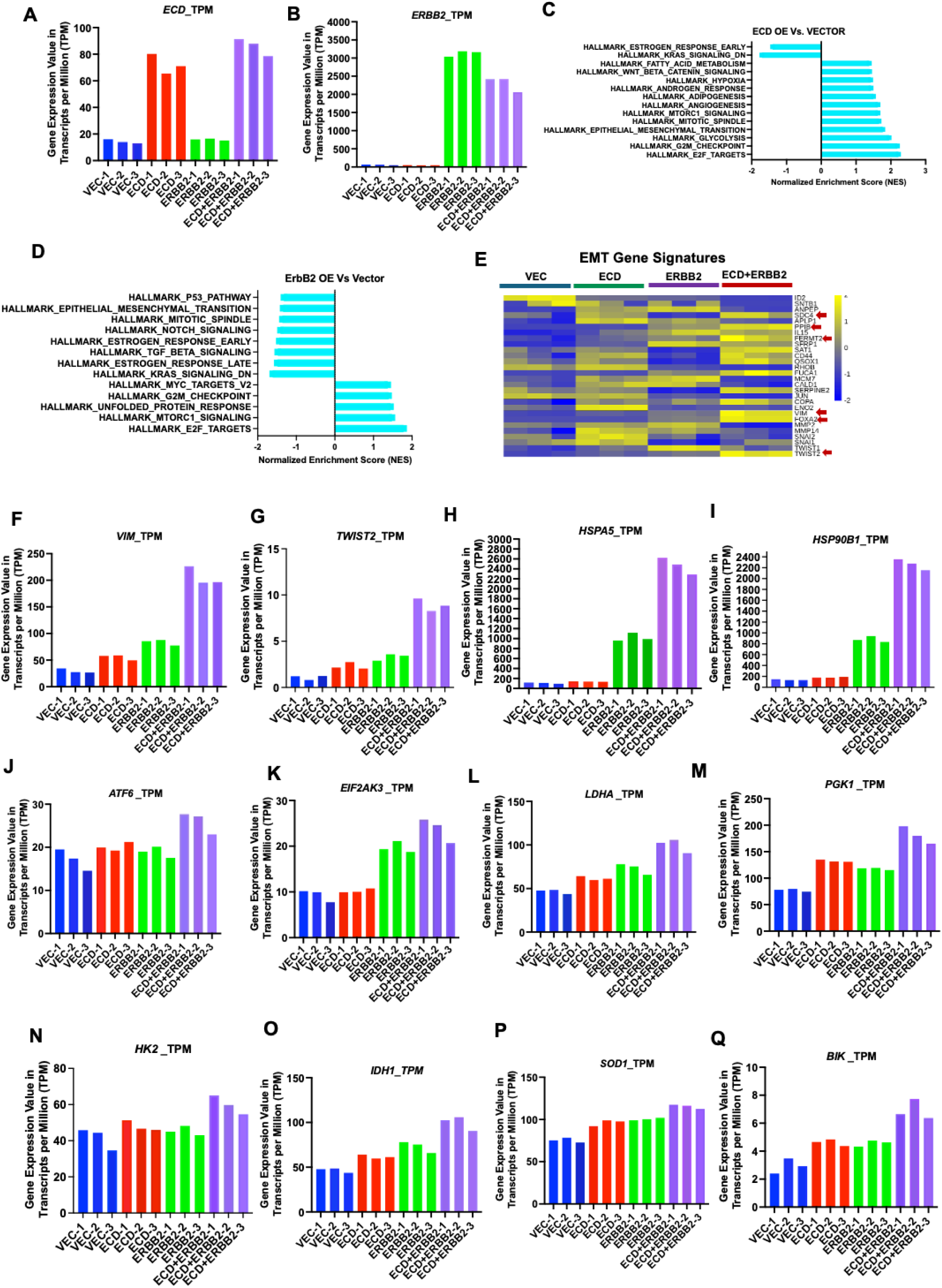
RNA-seq analyses comparison of vector, ECD overexpressing, ERBB2 overexpressing, and ECD+ERBB2 overexpressing 76NTERT cells and validation in 70NETRT cells. Transcripts per million (TPM) values for 76NTERT transductants RNA samples used for RNA-sequencing and analysis. Vector is shown in blue, ECD in red, ERBB2 in green, and ECD+ERBB2 in purple. *ECD* TPM (**A**) and *ERBB2* TPM (**B**) displayed for validation of samples used in analyses. (**C**) Bar graphs display enriched GSEA hallmark pathways including unfolded protein response (UPR), glycolysis, protein secretion, mTORC1 signaling, hypoxia signaling in ECD OE (overexpression) vs. Vector (**C**) and ERBB2 OE vs. Vector (**D**). X-axis represents normalized enrichment scores (NES) of the signaling pathways with significant nominal p-values (NOM p-val). Heatmap shows upregulated (yellow) and downregulated (blue) key EMT genes (**E**). TPM values of key EMT genes *VIM* **(F)** and *TWIST2* **(G)** are displayed. TPM values of key UPR-related genes *HSPA5* **(H),** *HSP90B1* **(I),** *ATF6* **(J),** *EIF2AK3* **(K)** and glycolytic genes *LDHA* **(L),** *PGK1* **(M),** *HK2* **(N),** *IDH1* **(O),** *SOD1* **(P),** *BIK1* **(Q)** are shown as bar graphs.

**Fig. S4.**
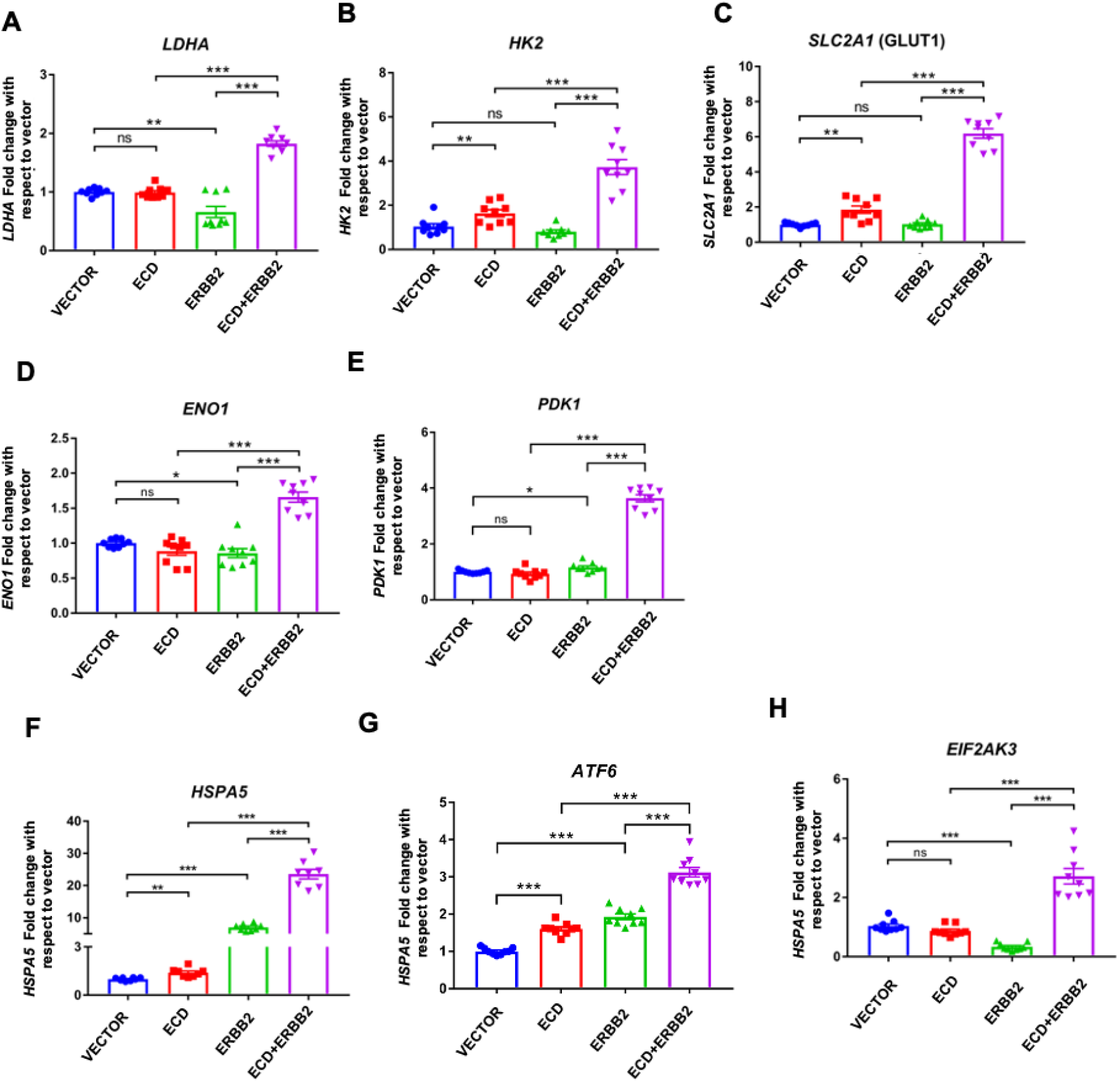
Confirmation of upregulated glycolytic genes and UPR-related genes in 70NETRT cells by qRT-PCR. Real-time qPCR analyses of glycolytic genes *LDHA* (**A**), *HK2* (**B**), *SLC2A1* (GLUT1) (**C**), *ENO1* (**D**) and *PDK1* **(E**) in 70NTERT transductants grown in serum-free DFCI-3 medium for 72 hours are presented. UPR related mRNA levels of *HSPA5 (*GRP78) (**F**), *ATF6* (**G**), *EIF2AK3* (PERK) (**H**) are presented. Relative levels of each transcript are expressed as fold change with respect to vector control cells after normalizing with the housekeeping gene, 18S and using the ΔΔCT method. Each bar graph indicates mean fold change +/− SEM from three experiments, each with three technical replicates (ns, p > 0.05, *, p<0.05, **, p < 0.01; ***, p < 0.001).

